# Unravelling the neurocognitive mechanisms underlying counterconditioning in humans

**DOI:** 10.1101/2024.07.29.605706

**Authors:** Lisa Wirz, Maxime C. Houtekamer, Jette de Vos, Joseph E. Dunsmoor, Judith R. Homberg, Marloes J.A.G. Henckens, Erno J. Hermans

**Author notes:** Corresponding author: Lisa Wirz, Kapittelweg 29, 6525 EN Nijmegen, The Netherlands. These authors contributed equally to this work (Lisa Wirz, Maxime C. Houtekamer, Judith R. Homberg, Marloes J.A.G. Henckens).

## Abstract

Counterconditioning (CC) aims to enhance extinction of threat memories by establishing new associations of opposite valence. While its underlying neurocognitive mechanisms remain largely unexplored, previous studies suggest qualitatively different mechanisms from regular extinction. In this functional MRI study, participants underwent categorical threat conditioning (CS+/CS-: images of animals/tools), followed by either CC (CS+ images reinforced with monetary rewards, n=24) or regular extinction (n=24). The following day, we assessed spontaneous recovery of threat responses and episodic memory for CS+ and CS-category exemplars. While the ventromedial prefrontal cortex (vmPFC) was activated during regular extinction, participants undergoing CC showed persistent CS+-specific deactivation of the vmPFC and hippocampus, and CS+-specific activation of the nucleus accumbens (NAcc). The following day, physiological threat responses returned in the regular extinction group, but not in the CC group. Counterconditioning furthermore strengthened episodic memory for CS+ exemplars presented during CC, and retroactively also for CS+ exemplars presented during the threat conditioning phase. Our findings confirm that CC leads to more persistent extinction of threat memories, as well as altered consolidation of the threat conditioning episode. Crucially, we show a qualitatively different activation pattern during CC versus regular extinction, with a shift away from the vmPFC and towards the NAcc.

## Introduction

Trauma-related disorders are prevalent and highly detrimental to the individual’s quality of life^1^. To treat these disorders, patients undergo exposure therapy in a safe therapeutic environment, causing threat responses to fade away^2^. Although exposure therapy may be successful initially, relapse often occurs and is the most prevalent remaining challenge in optimizing treatment efficacy. Research suggests that exposure therapy creates a safety memory that competes for expression with the original threat memory^3,4^, suggesting that relapse may occur because of relatively weak learning and retention of the safety memory. Therefore, identifying mechanisms that can be used to strengthen safety learning is a key step in advancing treatment for trauma-related disorders. A promising approach to strengthen safety learning is to create a new, positive association with the event that was previously linked to an aversive outcome. However, while there are indications that establishing positive associations can prevent relapse, the underlying mechanisms are poorly understood (for a review, see^5^).

To study threat responses in a controlled setting, aversive Pavlovian conditioning is typically used. A neutral stimulus (conditioned stimulus, CS; e.g., a picture) is coupled with a biologically aversive unconditioned stimulus (US; e.g., an electrical shock), after which the CS alone also elicits a conditioned threat response. Conditioned threat responses to the CS can be attenuated using extinction, during which the CS is repeatedly presented in absence of the US. However, early theories have suggested that threat responses may more easily be inhibited by engaging appetitive systems^6,7^. Indeed, experiments provide evidence that coupling a CS to a positive US after threat conditioning, a process known as aversive-to-appetitive counterconditioning (CC), may be superior to regular extinction. Specifically, CC compared to regular extinction was associated with a faster attenuation of learned threat responses^6,8^, stronger decreases in threat expectancy^9,10^, and more positive valence ratings of the CS^11–13^ immediately post-CC.

Tests for spontaneous recovery, reinstatement, and renewal can subsequently be used to evaluate the return of threat responses over time, after unsignaled presentation of the US, or in a novel context, respectively^3,14^. Thereby, one can investigate whether CC persistently attenuates threat responses. While early rodent studies showed that CC may be prone to the same relapse as extinction^15,16^, recent neurobiological work in rodents showed that CC can enhance the activation of an amygdala-striatal pathway, which is also recruited during extinction – albeit to a lesser degree –, and that CC compared to regular extinction can reduce the return of threat responses^17^. Recent studies suggest that CC may diminish the return of threat responses in humans as well. Specifically, it was shown that CC compared to regular extinction reduced renewal of previously learned food-allergy associations when presented in a novel context one day later^18^. Counterconditioning compared to regular extinction was also associated with reduced recovery of arousal and shock expectancy the following day^9,19^, as well as reduced reinstatement^9^.

Extinction learning appears to be mediated by activation of the ventromedial prefrontal cortex (vmPFC), which inhibits the expression of threat responses by suppressing amygdala activity^20–23^. When extinction is enhanced by replacing aversive with novel, neutral outcomes, the vmPFC was found to be engaged more effectively than during standard extinction^24^. When extinction is enhanced by replacing aversive outcomes with a reward (counterconditioning), evidence in rodents suggests stronger engagement of the ventral striatum, a region known to be involved in the anticipation and receipt of reward^25^. However, human studies only provide indirect evidence for such a mechanism since involvement of the ventral striatum could only be shown during spontaneous recovery^19^ or during reinstatement^26^, but not during CC itself. Although it was observed that brain areas of the fear network are reduced during CC versus regular extinction in humans^19^, it is unclear how this difference is achieved. Therefore, although evidence suggests that CC is more effective than regular extinction in preventing the return of threat responses the neural mechanisms are not well understood yet. It remains unclear whether CC is a form of enhanced extinction that is mediated by enhanced engagement of extinction networks, including the vmPFC, or whether it is driven by engagement of reward networks.

To investigate the qualitative differences between CC versus regular extinction further, category conditioning can be used, a procedure in which conditioned threat responses are learnt by coupling a US to conceptually linked exemplars that together form a category (e.g., pictures of animals)^27^. It allows for the typical measures of threat conditioning, but also provides the opportunity to probe episodic memory for the CS category exemplars^28^. Specifically, retrieval of a picture probes the retrieval of specific episodic elements, whereas retrieval of the category-threat association (fear retrieval) takes place at a conceptual level and may be semantic in nature. When episodic memory was probed 24h after CC and extinction, it was shown that memory for CS+ stimuli that had undergone CC was stronger than memory for CS+ stimuli that had undergone regular extinction^29^. This suggests that compared to regular extinction, CC can enhance episodic memory consolidation and potentially provide stronger retrieval competition against a threat memory.

To investigate the neural mechanisms that distinguish CC from regular extinction and to establish whether CC is indeed associated with a memory that is qualitatively different from the safety memory established during regular extinction, we performed a two-day fMRI study comparing CC versus regular extinction in a between-subjects design (**Figure 1A**). Participants underwent category conditioning and subsequently either aversive-to-appetitive CC (CC group) or regular extinction (Ext group; **Figure 1B-C**). During the CC task, participants in the CC group obtained monetary rewards depending on how quickly they responded to a cue superimposed on novel category exemplars from the CS+ category, a procedure similar to the monetary incentive delay (MID) task^30^. To maximize task similarity between tasks and groups, the cued-response element was kept consistent in all tasks (acquisition, CC/extinction, spontaneous recovery, reinstatement), but response-time contingent monetary rewards were only present during the CC task (**Figure 1F**). To assess the potential of CC versus regular extinction in persistently attenuating the expression of threat response, we tested retrieval of the threat memory and reinstatement of threat responses one day later (**Figure 1D**). Episodic memory for exemplars of the CS categories that were presented during threat conditioning and CC/extinction was assessed by means of a surprise memory test. To characterize pupil dilation responses (PDRs) and skin conductance responses (SCRs) during the anticipation of shock- and reward-reinforcement independently from prior conditioning, a separate valence-specific response characterization task was included at the end of the experiment (**Figure 1E**).

**Figure 1.**
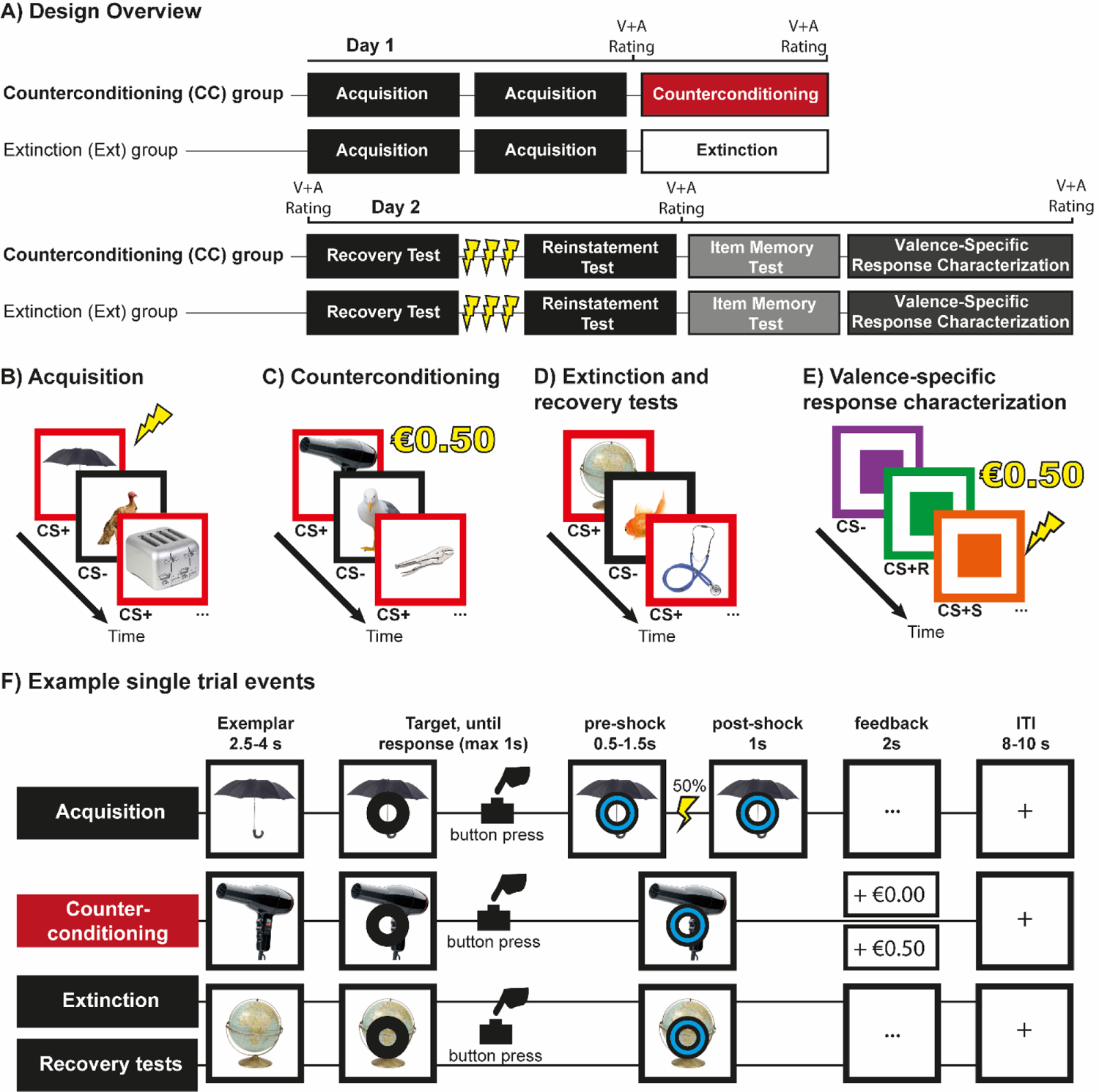
Overview of the experimental design. (A). Participants were assigned to the counterconditioning (CC) or extinction (Ext) group. On day 1, participants performed two blocks of acquisition of category-conditioned threat responses separated by a 30 second break, followed by CC or extinction. Day 2 consisted of a spontaneous recovery test, a reinstatement procedure and test, an item memory test and a valence-specific response characterization. Valence and arousal ratings for the different categories were taken before or after the tasks as indicated by ‘V+A Rating’. All tasks were performed in an MRI scanner. **(B)** During acquisition, participants viewed trial-unique exemplars of objects and animals. Exemplars of one category (CS+ animals or objects counterbalanced) were paired with a shock in 50% of trials. CS-trials were not reinforced. **(C)** Participants in the CC group could earn a monetary reward if they responded quickly enough to exemplars in the CS+ category. **(D)** Participants in the Ext group underwent extinction. During the extinction task, recovery test and reinstatement test, neither CS+ nor CS-exemplars were paired with a shock. **(E)** In the valence-specific response characterization task, participants viewed three different coloured squares. One colour was associated with shock (CS+S), one colour with reward (CS+R) and one colour served as CS-. The trial structure was otherwise identical to the acquisition and CC tasks. **(F)** In all Pavlovian tasks, trial onset was marked by presentation of a unique category exemplar. After a variable interval, a ring appeared, to which participants were instructed to respond as quickly as possible. Upon response, the ring shifted in colour as response confirmation. In the acquisition task, shocks could occur 0.5-1.5-seconds after the response window had elapsed (indicated as ‘pre-shock). The category exemplar and cue remained visible 1 second after potential shock administration (indicated as ‘post-shock’). During CC, participants received visual feedback for 2 seconds (+€0.50 approximately the fastest 70% of trials, +€0.00 on other trials). During the other tasks, participants viewed neutral feedback (three dots). Trials were separated by an 8-10 s intertrial interval, during which a fixation cross is displayed in the centre of the screen.

In line with previous results^9,19^, we hypothesized that CC compared to extinction would lead to a more persistent attenuation of threat responses. As indicated above, this could be mediated by two possible neural mechanisms: either through enhanced engagement of extinction networks, reflected by increased engagement of the vmPFC, or through a shift towards reward networks, reflected by activation of the ventral striatum. Based on previous results^29^, we expected stronger episodic memory for CS exemplars presented during CC, whereas regular extinction would not show such a strengthening effect.

## Results

In the valence-specific response characterization task (Figure 1E), we observed that both threat and reward-anticipation induced strong arousal-related PDRs and SCRs (see **Supplementary Information**). However, PDRs allowed for a better differentiation of the two compared to the CS-(**Supplementary Figure 1A**). Therefore, we focused on PDRs in all analyses and refer to the Supplementary Information for details on the analysis of SCRs. During the acquisition task, both groups showed comparable and successful acquisition of differential conditioned threat responses (PDR means ± SD: CC CS+=1.085±.030, CC CS-=1.054±.033, Ext CS+=1.084±.050, Ext CS-=1.050±.035; for PDR, SCR and fMRI results see **Supplementary Information**).

### Extinction and aversive-to-appetitive counterconditioning

After threat acquisition, participants in the CC group underwent CC, while participants in the Ext group underwent regular extinction (see Figure 1A for design overview). Across both groups and phases (early vs. late), we observed retention of conditioned differential PDRs (CS-type (CS+, CS-) x Phase (Early, Late) x Group (CC, Ext) rmANOVA, main effect CS-type: F(_1,34_)=15.393, p<0.001, η²=0.312, Figure 2A), as well as a decrease in PDRs over the course of the task (main effect phase: F(_1,34_)=10.121, p=0.003, η²=0.229). These findings are in contrast to our expectation of a CS-type x Phase x Group interaction. Specifically, we expected differential PDRs to become extinguished in the Ext group, while being sustained in the CC group, potentially due to increased reward anticipation. Extinction in the Ext group however already occurred during the early phase (paired t-test, early CS+ vs. CS-, p=0.233), and differential responses did not change towards the late phase (p=0.979). As a result, we found distinct differential conditioned PDRs throughout the CC/extinction task between groups (CS-type x Group interaction: F(_1,34_)=6.053, p=0.019, η²=0.151), with participants undergoing CC showing stronger PDRs to CS+ vs. CS-category exemplars (paired t-test average CS+ vs. CS-, t(20)=3.602, p=0.002, CS+: 1.07±0.04, CS-: 1.04±0.04), whereas differential PDRs were extinguished in participants undergoing extinction (paired t-test average CS+ vs. CS-, p=0.246, CS+: 1.05±0.04, CS-: 1.04±0.04). Results of the valence-specific response characterization task showed that differential PDRs can also be indicative of anticipation of reward (**Supplementary Figure 1A**). Thus, while PDRs in the Ext group indicated that differential conditioned threat responses were successfully extinguished, differential PDRs persisted in the CC group, likely reflecting reward anticipation. Differential SCRs persisted during the late phase of both CC and extinction but were no longer detectable in the last two trials and were comparable between groups (see **Supplementary Information**).

**Figure 2.**
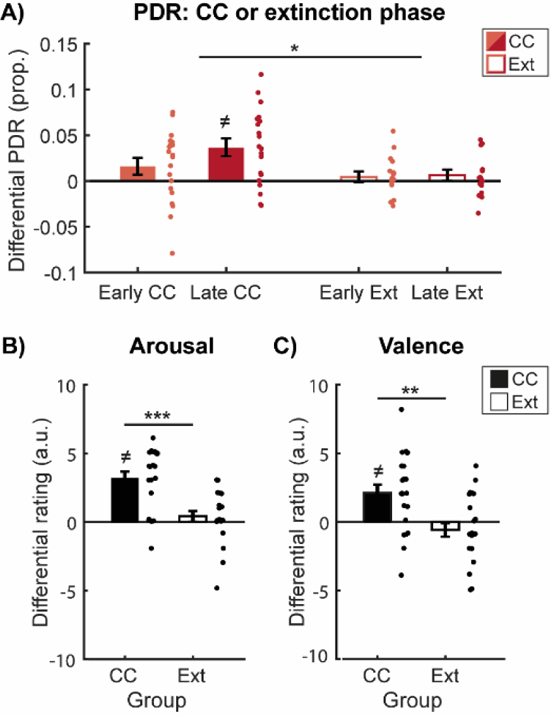
Differential PDRs during CC/extinction and explicit ratings of arousal and valence provided after the counterconditioning or extinction phase. (A) Differential PDRs for the early (light red) and late (dark red) phase of counterconditioning (CC, solid bars) or extinction (EXT, open bars). Participants undergoing CC showed increased differential PDRs as compared to participants undergoing extinction. (B) Arousal and (C) valence ratings displayed separately for participants assigned to the counterconditioning (CC, solid bars) and extinction (EXT, open bars) groups. Participants that had undergone CC gave stronger differential arousal scorings than participants that had undergone extinction. In addition, participants that underwent CC showed flipped differential valence ratings: while valence differential valence ratings were negative after extinction, the direction reversed to positive differential ratings after CC. Error bars represent ± standard error of the mean. *=p<0.05, **=p<0.01, ***=p<0.001, ≠ indicates that the bar is significantly different from 0.

Valence and arousal ratings provide further support for the extinction of differential responses in the Ext group and positive, reward-induced arousal for CS+ items in the CC group (Figure 2B-C). Differential valence ratings for the CS+ and CS-differed between groups after the CC/extinction task (CS-type (CS+, CS-) x Group (CC, Ext) rmANOVA, CS-type x Group interaction: F(_1,44_)=12.054, p=0.001, η²=0.215). Participants in the CC group rated CS+ stimuli more positive than CS-stimuli (t(21)=3.469, p=0.002, CS+: 7.5±0.30, CS-: 5.41±0.38), while participants in the Ext group gave both categories similar valence ratings (p=0.245, CS+: 5.63±0.32, CS-: 6.21±0.28). Since there were group-dependent differences in valence ratings after acquisition (the CS-category was rated more positively by the Ext compared to the CC group, see **Supplementary Information**), we ran an additional analysis, adding differential valence ratings after fear acquisition as covariate. Results suggested that the group difference in differential valence ratings after CC/extinction remain (main effect Group: F_(1,43)_=7.364, p=0.010, η²=0.146). Differential arousal ratings for the CS+ and CS-also differed between groups (CS-type (CS+, CS-) x Group (CC, Ext) rmANOVA, CS-type x group interaction: (F(_1,44_)=20.862, p<0.001, η²=0.322). Participants in the CC group reported higher arousal levels for the CS+ category than for the CS-category (t(21)=6.370, p<0.001, CS+: 6.64±0.20, CS-: 3.45±0.38) while participants in the Ext group gave similar arousal ratings for the CS+ and CS-categories (p=0.290, CS+: 4.21±0.43, CS-: 3.80±0.40). Taken together, more positive valence and higher arousal ratings for the CS+ in the CC group as compared to the Ext group further support the interpretation of increased differential PDRs reflecting arousal induced by reward anticipation.

### CC prevents differential spontaneous recovery

To investigate whether CC prevented the spontaneous recovery of differential conditioned threat responses one day later (see Figure 1A for design overview), we compared PDRs in the last two trials of the CC/extinction task and the first two trials of the spontaneous recovery test in a CS-type (CS+, CS-) x Group (CC, Ext) x Phase (last two trials of CC/extinction, first two trials of the spontaneous recovery test) rmANOVA [16 participants were excluded due to (partially) missing data, leaving 15 participants in both groups (total N=30)]. We expected the Ext group to show an increase in PDRs from the extinction task to the spontaneous recovery task, while we expected PDRs for the CC group to remain stable or decrease. Critically, differential spontaneous recovery of PDRs differed between groups (Group x CS-type x Phase interaction: F(_1,28_)=6.329, p<0.018, η²=0.184, Figure 3). While the CC group showed a decrease in differential PDRs from CC to spontaneous recovery (t(14)=-1.807, p=0.046, one-tailed, CC: 0.34±0.2, spontaneous recovery: −0.01±0.18), the Ext group showed an increase in differential PDRs (t(14)=1.850, p=0.043, one-tailed significance, extinction: 0.11±0.01, spontaneous recovery: 0.04±0.02). To conclude, while we observed differential spontaneous recovery in the Ext group, we did not find evidence for differential spontaneous recovery in the CC group, suggesting that CC attenuated the recovery of threat-responses compared to regular extinction.

**Figure 3.**
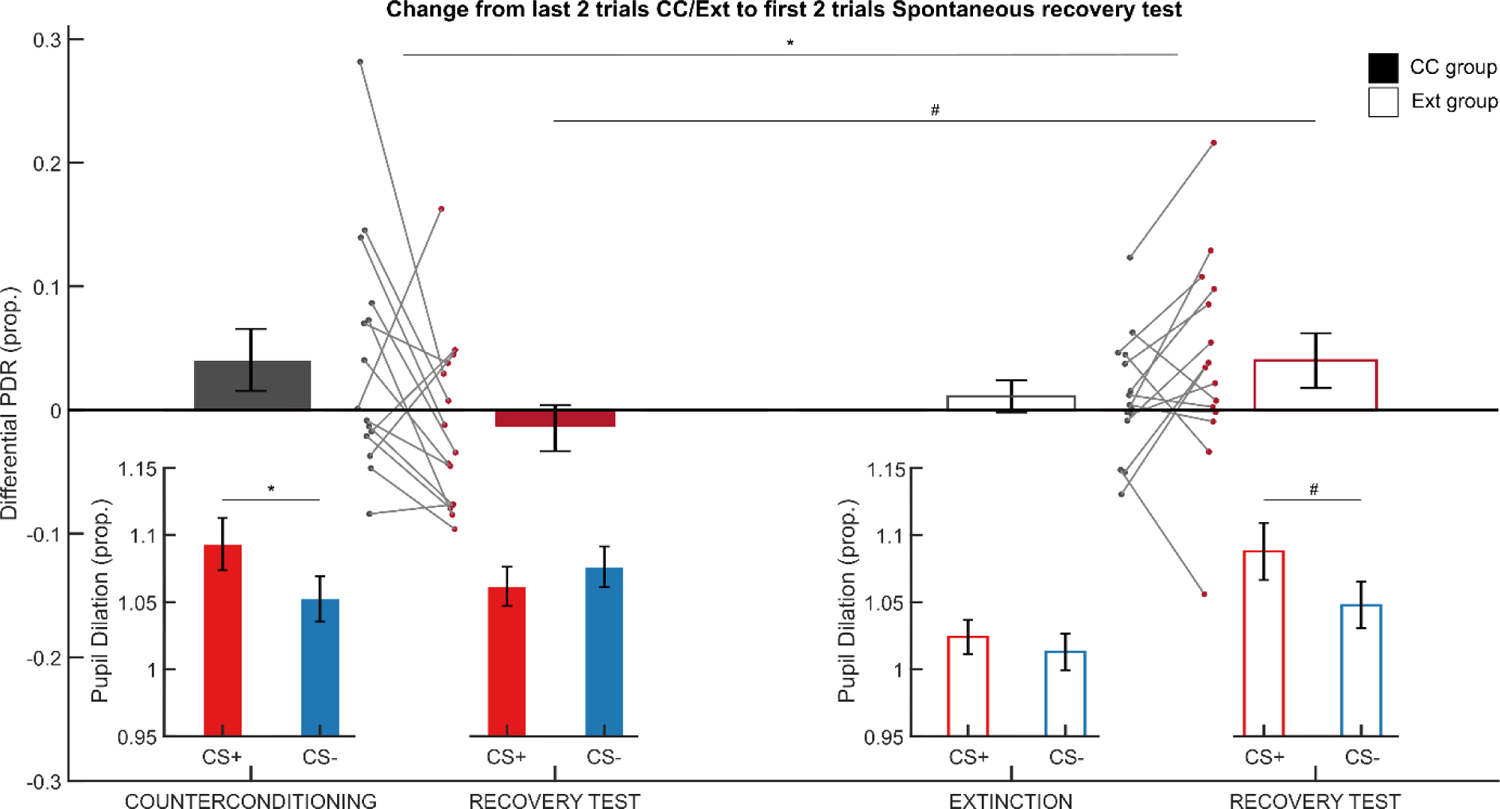
Differential PDRs during the last two trials of counterconditioning or extinction and the first two trials of the spontaneous recovery test. Differential PDRs show selective spontaneous recovery after extinction (Ext, open bars) but not after counterconditioning (CC, solid bars). During the first two trials of the spontaneous recovery test, differential PDRs are increased in the Ext group as compared to the CC group. Insets show PDRs to the CS+ (red) and CS-(blue) during the last two trials of CC/Ext and the first two trials of the spontaneous recovery test. While the Ext group shows differential responding during the spontaneous recovery test, the CC group does not. Error bars represent ± standard error of the mean. *=p<0.05, #=p<0.05 one-tailed significance.

However, since participants undergoing CC showed persistent differential PDRs during the last two trials of the CC phase (likely due to reward anticipation), while participants undergoing extinction did not, we additionally explored whether there was differential responding during the first two trials of the spontaneous recovery test. During the first two trials of the spontaneous recovery test, participants in the CC group showed decreased differential PDRs as compared to the Ext group (CS-type (CS+, CS-) x Group (CC, Ext) rmANOVA, CS-type x Group interaction: F(_1,29_)=3.901, p=0.029, one-tailed, η²=0.119).

Further exploration within the groups confirmed that participants in the CC group did not show retention of differential responses (paired t-test, CS+ and CS-responses during the first two trials of the spontaneous recovery test, p=0.219, one-tailed), while the Ext group did show increased responses to the CS+ as compared to the CS-(t(14)=1.958, p=0.035, one-tailed). Thus, both the differential spontaneous recovery of PDRs between sessions, and differential responding within the first two trials of the spontaneous recovery test suggested that CC prevented spontaneous recovery of differential responses compared to extinction. SCRs did not show differential recovery and were comparable between groups (see **Supplementary Information**).

CC also appeared to have lasting beneficial effects on valence ratings compared to extinction. At the start of the second testing day, differential valence ratings continued to differ between groups (CS-type (CS+, CS-) x Group (CC, Ext) rmANOVA, CS-type x Group interaction: F(_1,44_)=5.160, p=0.028, η²=0.105). While participants in the CC group gave similar valence ratings to both categories (p=0.179, CS+: 6.3±0.34, CS-: 5.4±0.35), participants in the Ext group gave more negative valence ratings to the CS+ category than to the CS-category (t(23)=-1.964, p=0.031 one-tailed test, CS+: 5.5±0.30, CS-: 6.3±0.24), also illustrative of relapse of threat associations. Since the CC group received no rewards during the spontaneous recovery task and was aware of this, it was expected that the effect of reward anticipation on PDRs would be weakened in the CC group. So while PDRs did not differ between CS+ and CS-in the CC group because no rewards were given, CS+ and CS-items were still rated of similar valence. In comparison, the Ext group rated the CS+ significantly more negative and threat responses to the CS+ returned.

While participants in the CC group showed heightened differential arousal ratings immediately after CC as compared to ratings from participants who had undergone extinction (Figure 2B), participants in both groups gave comparable differential arousal ratings at the start of the second day immediately before the spontaneous recovery test (CS-type (CS+, CS-) x Group (CC, Ext) rmANOVA, main effect of CS-type: F(_1,44_)=10.932, p=0.002, η²=0.022, CS+: 4.8±0.28, CS-: 3.9±0.24). Likewise, response times to the CS+ and CS-during the first two trials of the spontaneous recovery task were similar across both groups (all p’s>0.2). These findings may suggest that differential arousal evoked by the categories was similar in both groups immediately before and during the spontaneous recovery test.

The spontaneous recovery test was followed by a reinstatement procedure, consisting of three unsignaled shocks, and a reinstatement test. However, mean PDRs decreased from spontaneous recovery to reinstatement (t(30)=3.063, p=0.005, last two trials of spontaneous recovery: 1.04±0.01, first two trials of reinstatement: 1.01±0.01). Given that we did not observe successful reinstatement in either group, our reinstatement test was not informative on whether CC can lead to a more persistent attenuation of threat responses as compared to regular extinction. A full description of PDR and SCR results of the reinstatement test can be found in the **Supplementary Information**.

### Distinct CS-type specific activation for extinction and appetitive counterconditioning

During the CC/extinction task, whole brain analysis revealed that CS-type specific activation changed differentially between the two groups in a large cluster encompassing multiple regions in the medial temporal lobe (Group x CS-type x Phase interaction, cluster size = 1760 mm^3^, p=0.034, whole-brain FWE-corrected, Figure 4B and **Table 1**). We further investigated the anatomical location of the cluster using our ROIs to probe for activity and found that the effect encompassed the amygdala. To further investigate the interaction effect in the amygdala, we extracted parameter estimates from the complete bilateral amygdalae (Automated Anatomic Labelling, AAL, atlas in the WFU PickAtlas toolbox in MN152 space) and performed post-hoc comparisons. In the early phase, CS-type specific responses differed between the groups (t(1,44)=2.173, p=0.035, CC: 0.18±0.08, Ext: −0.073±0.08). Specifically, the CC group showed increased amygdala activation to the CS+ as compared to the CS-(t(23)=2.210, p=0.037) while that was not the case in the Ext group (p=0.390). In the late phase, differential responses were comparable between the groups (p=0.503).

**Figure 4.**
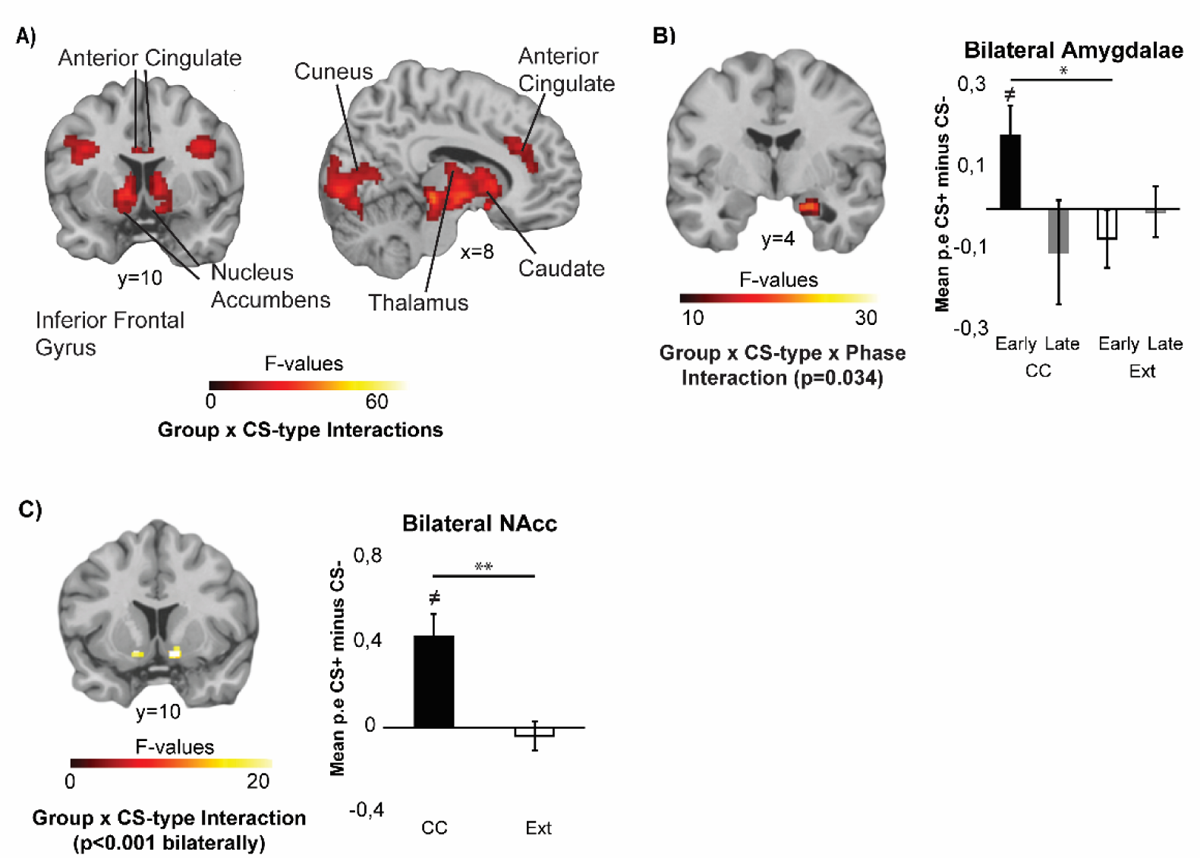
Stimulus-type specific activation differs between participants undergoing CC versus extinction. **A**. Whole-brain Group x CS-type interaction effects revealed distinct stimulus-specific activation of regions including the anterior cingulate, cuneus, nucleus accumbens, caudate, thalamus and inferior frontal gyrus during the counterconditioning vs. extinction phase. Panel A displays group F-images (see **Table 1** for directions) FWE-corrected at p<0.05, cluster-forming threshold p=0.001. **B**. The right amygdala showed a Group x CS-type x Phase interaction during the CC/extinction task, indicating that CC compared to extinction is associated with decreased activation of the amygdala**. C.** The bilateral NAcc showed a Group x CS-type interaction during the CC/extinction task, revealing increased NAcc activation in response to the CS+ compared to the CS-in the CC but not in the Ext group. Panel B and C display group F-images FWE-SVC at p<0.05, cluster-forming threshold p=0.001, along with post-hoc tests on mean parameter estimates from the complete ROI included in the analyses. ** p<0.01, * p<0.05, ≠ indicates that the value is significantly different from 0.

**Table 1.** Whole-brain main effects of group (CC, Ext), CS type (CS+, CS-) and phase (early, late) and interactions, during the counterconditioning/extinction task. Cluster-forming threshold p=0.001, FWE-corrected at p<0.05, clusters were labelled using the Talairach Daemon atlas and the AAL atlas for ROIs. For each cluster, the peak voxel coordinates (MNI space) and regions are reported, and additional regions contained within the cluster are added in italics. See **Supplementary Table 1** for main effects of CS-type.

| Peak MNI coordinate |  |  |  |  |  |  |  |  |
| --- | --- | --- | --- | --- | --- | --- | --- | --- |
| Region | Cluster | x | y | z | Size (mm3) | pFWE (cluster) | Peak F-value | Direction |
| Group x CS-type x phase |  |  |  |  |  |  |  |  |
| Parahippocampal Gyrus BA34R<br>Parahippocampal Gyrus Amygdala, Uncus BA34R | 1 | 18 | -8 | -20 | 1760 | 0.034 | 23.40 | CS+>CS- difference increases from early to late phase for CC, not for Ext |
| Group x CS-type |  |  |  |  |  |  |  |  |
| Lateral Geniculum Body LR, Caudate Head LR, Thalamus LR, Lentiform Nucleus LR | 1 | 2 | -26 | -18 | 29920 | <0.001 | 73.15 | (CC CS+>CS-) > (Ext CS+>CS-) |
| Cuneus L<br>Lingual Gyrus BA17/BA18 LR, Posterior Cingulate LR, Cuneus BA18R, Cuneus BA30L Declive R | 2 | -6 | -96 | 2 | 23272 | <0.001 | 43.50 |  |
| Inferior Frontal Gyrus BA47L<br>Insula BA13 L | 3 | -36 | 18 | -6 | 4504 | 0.009 | 30.62 |  |
| Extra-Nucleus R | 4 | 30 | 26 | 2 | 3136 | 0.016 | 37.67 |  |
| Superior Temporal Gyrus L<br>Superior Temporal Gyrus BA41 L, Transverse Temporal Gyrus L | 5 | -60 | -44 | 14 | 9088 | 0.002 | 43.56 |  |
| Transverse Temporal Gyrus BA41 R<br>Superior Temporal Gyrus R, Superior Temporal Gyrus BA42/BA22R | 6 | 44 | -22 | 12 | 7784 | 0.003 | 42.17 |  |
| Anterior Cingulate BA32R<br>Anterior Cingulate BA32L, Cingulate Gyrus R | 7 | 6 | 30 | 26 | 8880 | 0.002 | 27.90 |  |
| Precentral Gyrus L<br>Inferior Frontal Gyrus L | 8 | -36 | 0 | 30 | 3624 | 0.014 | 30.10 |  |
| Precentral Gyrus R<br>Sub-Gyral R | 9 | 40 | 2 | 32 | 4056 | 0.011 | 40.64 |  |
| Precentral Gyrus BA6L<br>Middle Frontal Gyrus BA6L | 10 | -44 | -6 | 52 | 2184 | 0.028 | 24.34 |  |
| Angular Gyrus R<br>Supramarginal Gyrus R | 11 | 54 | -60 | 36 | 1944 | 0.032 | 24.18 |  |
| Group x Phase |  |  |  |  |  |  |  |  |
| No significant clusters |  |  |  |  |  |  |  |  |
| CS-type x Phase |  |  |  |  |  |  |  |  |
| No significant clusters |  |  |  |  |  |  |  |  |
| Group |  |  |  |  |  |  |  |  |
| No significant clusters |  |  |  |  |  |  |  |  |
| Phase |  |  |  |  |  |  |  |  |
| Inferior Frontal Gyrus R<br>Inferior Frontal Gyrus BA45 R | 1 | 30 | 26 | 8 | 4848 | 0.006 | 40.27 | Early Phase > Late Phase |
| Insula L<br>Superior Temporal Gyrus BA22, Precentral Gyrus L | 2 | -28 | 26 | 0 | 4368 | 0.007 | 38.41 |  |
| Postcentral Gyrus L | 3 | -54 | -24 | 22 | 1768 | 0.031 | 23.75 |  |

Whole-brain analysis further revealed a number of clusters showing distinct CS-specific activations between groups throughout the task, including the anterior cingulate, cuneus, nucleus accumbens, caudate, thalamus and inferior frontal gyrus (Figure 4A, **Table 1**). The group and stimulus-specific activation of the NAcc was in line with a priori expectations for the CC phase (Figure 4C). To further explore this effect, averaged parameter estimates from the bilateral NAcc ROI (mask acquired from the IBASPM 71 atlas in the WFU PickAtlas toolbox in MNI152 space) were extracted. Across the bilateral NAcc, differential activation was increased in the CC as compared to the Ext group (t(44)=2.731, p=0.009, CC: 0.37±0.10, Ext: 0.04±0.06), with the CC showing increased NAcc activation to the CS+ compared to the CS-(t(23)=6.194, p<0.001, CS+: 0.59±1.12, CS-: 0.16±0.09) whereas the Ext group did not (p=0.574).

Contrast estimates in further a priori defined ROIs during the CC/extinction task were submitted to a Group (CC, Ext) x CS-type (CS+, CS-) x Phase (early, late) rmANOVA (Figure 5). The bilateral hippocampi (right hippocampus cluster size: 664 mm^3^, p=0.001, FWE-SVC, left hippocampus cluster size: 112 mm^3^, p=0.024, FWE-SVC) and the left vmPFC (mask defined as bilateral gyrus rectus and medial orbital gyri, cluster size = 160 mm^3^, p=0.013, FWE-SVC) showed differentially changing CS-type-specific activations between the groups (Group x CS-type x Phase interaction). While CS+-specific suppression of these regions appeared to increase during the CC task, this was not the case during the extinction task. Post-hoc comparisons on averaged parameter estimates in the bilateral hippocampi confirmed that stimulus-specific suppression increased during the course of the task in the CC group (t(23)=3.280, p=0.003, early CS+-CS-: 0.054±0.07, late: −0.150±0.07), but not in the Ext group (p=0.266). Post-hoc comparisons across the vmPFC ROI also revealed increased CS+-specific suppression in the CC group compared to the Ext group (t(44)=2.221, p=0.032, CC: −0.189±0.06, Ext: −0.070±0.10). While the extinction group showed increased CS+-specific activation from the early to the late phase of the extinction task (t(21)=2.235, p=0.036, early CS+: −0.149±0.08, late CS+: 0.040±0.09), the CC group did not (p=0.120). During the late phase, the CC group showed increased vmPFC deactivation to the CS+ compared to the CS-(t(23)=3.174, p=0.004, late CS+: −0.284±0.06, late CS-: −0.095±0.05), while the Ext group did not (p=0.503). Thus, across both the hippocampus and the vmPFC, CC induced increased stimulus-specific suppression. During the spontaneous recovery task, a priori defined regions of interest did not reveal any effects (see **Supplementary Information**).

**Figure 5.**
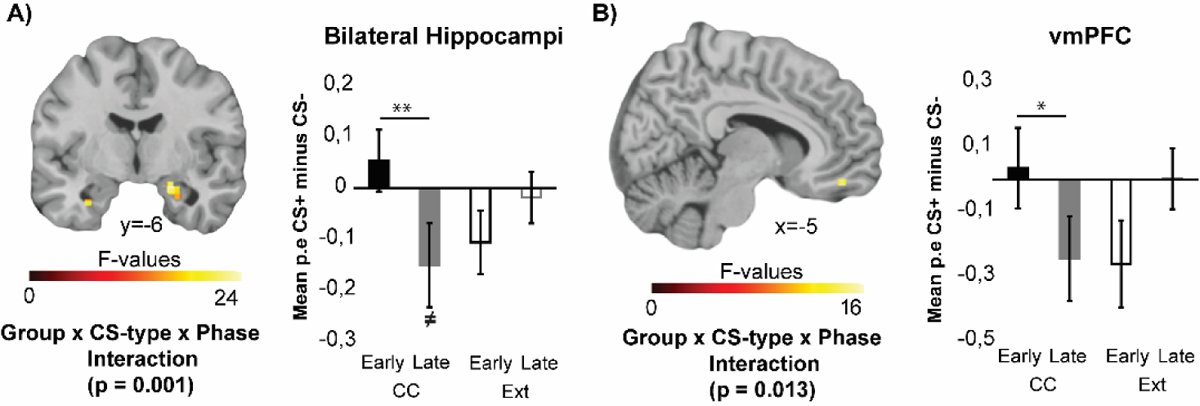
ROI analyses during the CC/extinction task reveal distinct activity in the hippocampus and left vmPFC. During the CC/extinction task, stimulus-specific activation of the hippocampus (C) and left vmPFC (**D**) changes differently between groups. ** p<0.01, * p<0.05, ≠ indicates that the bar is significantly different from 0. Group F-images FWE-SVC at p<0.05, cluster-forming threshold p=0.001, along with post-hoc tests on mean parameter estimates from complete ROI included in the analyses.

### Counterconditioning retroactively enhances item recognition for conditioned exemplars

Following the reinstatement test, participants completed a surprise item recognition test approximately 24 hours after acquisition and the CC/extinction task. One outlier was excluded from this analysis (CS-false alarm rate = 0.91). Threat conditioning has previously been shown to enhance 24-hour item recognition for category exemplars presented during the acquisition phase^27^. However, this enhancement for CS+ items did not extend to items presented during an extinction session separated from the acquisition phase by a short break^31^. We therefore analysed item recognition for the CS+ and CS-during acquisition and the CC/extinction phase separately to examine whether the groups differed in recognition memory performance (Figure 6).

**Figure 6.**
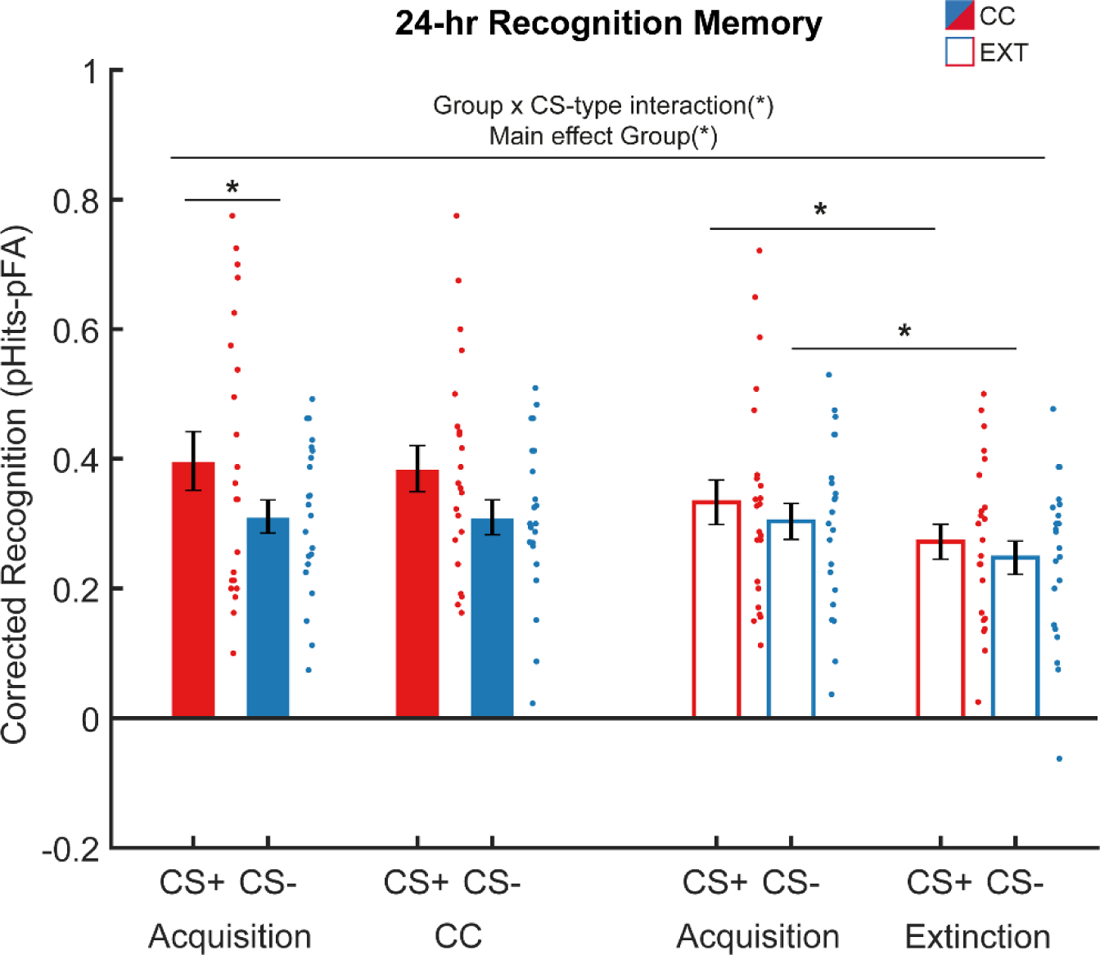
Twenty-four hour recognition memory results. During acquisition and extinction on the first day of the experiment, participants viewed trial-unique exemplars from two semantic categories (objects, animals) that served as CS+ and CS-. The next day, participants completed a surprise memory test for these items, mixed with an equal number of novel exemplars. Participants recognized relatively more items from the CS+ category, and participants that underwent CC showed improved item recognition compared to participants in the Ext group. Error bars represent ± standard error of the mean. *=p<0.05.

Corrected recognition scores (hits probability-false alarms probability) were subjected to a task (acquisition, CC/extinction task) x CS-type (CS+, CS-) x Group (CC, Ext) rmANOVA, including CS+-category (animals, tools) as covariate. Overall, participants showed better memory for items from the CS+ category (main effect of CS-type: (F(_1,42_)=10.615, p=0.002, η²=0.202) and participants who underwent CC showed better memory as compared to participants who underwent extinction (main effect of Group: (F(_1,42_)=4.963, p=0.031, η²=0.106). Stimulus-type specific item recognition differed between the CC and Ext groups (CS-type x Group interaction: F(_1,42_)=4.535, p=0.039, η²=0.094). While participants in the CC group showed better recognition memory for the CS+ category compared to the CS-category (t(22)=2.531, p=0.019, means ± SD: CS+ 0.39±0.17, CS-0.31±0.10), this was not the case for participants in the Ext group (t(23)=0.889, p=0.384, means ± SD: CS+ 0.30±0.13, CS-0.28±0.11). Although the effect of stimulus-type was stronger for tools as CS+, this was not different between groups (see **Supplementary Information**). Thus, across the acquisition and CC/extinction phase, participants who underwent CC showed a stronger enhancement of CS+ memory compared to the participants that underwent extinction.

To further investigate to what extent CC retroactively affected memory for items presented during the acquisition task, we examined item recognition during acquisition and the CC/extinction tasks separately. While threat conditioning increased memory for CS+ items presented during the acquisition task across both groups (main effect CS-type: (F(_1,42_)=18.147, p=<0.001, η²=0.302), subsequent CC enhanced this effect (Group x CS-type interaction: (F(_1,42_)=5.112, p=0.029, η²=0.109). Post-hoc tests revealed increased item memory for the CS+ category compared to the CS-category presented during acquisition in the CC group (t(22)=2.341, p=0.029, means ± SD: CS+ 0.40±0.21, CS-0.31±0.12) but not in the Ext group (t(23)=0.818, p=0.422, means ± SD: CS+ 0.33±0.16, CS-0.30±0.13).

Again, although the effect of stimulus-type was stronger for tools as CS+, this was not different between groups (see **Supplementary Information**). As the acquisition task was identical between groups, it appears that CC in comparison to extinction retroactively enhanced memory for CS+ items. For items presented during the CC/extinction task, overall item recognition was better in the CC group compared to the Ext group (main effect group: F(_1,42_)=8.706, p=0.005, η²=0.172, means ± SD: CC 0.35±0.12, Ext 0.26±0.09). Thus, compared to regular extinction, CC enhanced recognition of items presented during CC, but interestingly also strengthened the emotional memory enhancement of CS+ exemplars presented during acquisition, suggesting that immediate CC may alter consolidation of a prior threat learning episode.

Following previous work^29,31,32^, we explored stimulus-type specific decreases in item recognition between tasks, as well as within-phase differences between item recognition for the CS+ and CS-within each group. As expected, a post-hoc paired samples t-test showed that participants in the Ext group remembered significantly more CS+ items from the acquisition phase as compared to the extinction phase (t(23)=2.238, p=0.036, means ± SD: acquisition 0.33±0.16, extinction 0.27±0.13). In contrast, participants who had undergone CC remembered CS+ items presented during acquisition and CC equally well (t(22)=0.390, p=0.701, means ± SD: acquisition 0.40±0.21, CC 0.38±0.16). Thus, while recognition memory for items encoded during the extinction task was substantially weaker than memory for items from the acquisition task, this was not the case for items presented during CC.

## Discussion

This study aimed to test whether CC compared to regular extinction can lead to a more persistent attenuation of threat responses, and to investigate whether this is mediated by neural mechanisms reflecting extinction-related enhanced engagement of the vmPFC or engagement of reward-focused networks. We found that CC prevented differential spontaneous recovery of PDRs compared to regular extinction, suggesting that CC reduces the recovery of threat responses. Notably, there are individual differences (some participants in both groups show the opposite pattern), which should be further investigated in future studies with larger sample sizes, as it is crucial to identify who will respond to treatments based on the principles of standard extinction or counterconditioning. Our fMRI results suggested that CC engages different neural mechanisms compared to extinction. Most notably, while the extinction group showed an increase in CS+-specific vmPFC activation during extinction, the CC group showed CS+-specific deactivation of the vmPFC that persisted throughout the late phase of CC. Furthermore, CC led to increased NAcc activation for the CS+ compared to the CS-, whereas this was not the case for extinction. Lastly, phase- and stimulus-specific activation of the hippocampus and the amygdala differed between extinction and CC. Compared to extinction, CC led to increased activation of the amygdala in the early phase, and increasing stimulus-specific deactivation of the hippocampus over the course of the early and late phases. In addition, CC retroactively enhanced item recognition for conditioned exemplars presented during acquisition and strengthened memory for conditioned exemplars presented during CC compared to extinction.

The mechanism underlying CC appears to be qualitatively different from the mechanism underlying regular extinction. Regular extinction, which has been interpreted as an implicit form of emotion regulation^33^, is associated with activation of the vmPFC^23,34^, which is thought to inhibit the expression of threat responses by suppressing amygdala activity^20–23^. In comparison to regular extinction, novelty facilitated extinction, a form of enhanced extinction, in which aversive events are replaced with novel neutral outcomes, has shown stronger CS+-specific vmPFC activation^24^. If CC was similarly mediated by enhanced recruitment of extinction networks, we would have expected increased activation of the vmPFC, yet we observed a CS+-specific deactivation of the vmPFC during CC, disproving this hypothesis. Interestingly, deactivation of the vmPFC during CC was also found in studies investigating a form of counterconditioning induced by means of real-time fMRI decoded neurofeedback^35,36^. During neurofeedback CC, participants implicitly learned to obtain monetary rewards by generating a representation of the target CS+ in the visual cortex^35^. After neurofeedback CC, reductions in threat responses were stronger in participants showing stronger vmPFC deactivation, suggesting that vmPFC disengagement may be associated with fear reductions^35^. Taken together, both our findings and previous neurofeedback studies suggest that in contrast to enhanced extinction, CC disengages the vmPFC. Given that we replicate this finding using a different approach that includes direct exposure to the CS+, vmPFC disengagement may be a distinguishing characteristic of CC, which could indicate that CC, in contrast to regular extinction, does not involve an implicit regulation strategy. The observed pattern of activity, including vmPFC deactivation further bears resemblance to activity patterns observed during goal-directed eye movements in an experimental model of eye-movement desensitization and reprocessing (EMDR), which has also been shown to improve extinction learning^37^. A similar activity pattern and effect has also been found for working memory-like tasks, such as a game of Tetris^38–40^. Given that the above-mentioned tasks associated with vmPFC deactivation share their strong engagement of working memory and/or endogenous attention mechanisms, thereby engaging the executive control-network, deactivation of the vmPFC and hippocampus could be the result of a deactivated default mode network due to competition between activation of large scale brain networks^41–43^.

The CC procedure led to clear CS+-specific activation of the NAcc, which is in line with expectations for reward anticipation in tasks with a monetary incentive delay aspect^44^. Activation of the ventral striatum has also been reported for active avoidance, and may be generally associated with instrumental actions as opposed to passive delivery of an outcome^45,46^. In line with studies on active avoidance, delivery of a reward contingent on instrumental actions has been shown to yield CC that is more resistant to renewal^47^. CS+-specific activation of the NAcc was not seen in participants undergoing extinction, suggesting that this activation is specific to CC. However, previous work in rodents revealed an amygdala-ventral striatum (NAcc) pathway that is activated during extinction training^17^. The recruitment of this pathway was shown to be enhanced during CC, and reduced the return of fear^17^, suggesting that CC may in fact enhance activation of reward-related networks that are weakly activated by extinction. Indeed, fMRI studies in humans that modelled prediction error for omitted aversive outcomes during extinction training (i.e. outcomes “better-than-expected”) showed involvement of the NAcc^48–50^. Possibly, activation of the NAcc during extinction is limited to early extinction trials generating prediction errors. Nevertheless, based on our findings, it appears that sustained CS+-specific activation of the NAcc is a distinct mechanism underlying CC but not extinction, which is potentially associated with instrumental actions. In comparison to more implicit forms of emotion regulation as is the case in regular extinction, CC may thus be a more active coping strategy, which is more effective in persistently preventing the return of threat responses^45^.

A recent neuroimaging study suggests that the neural differences between regular extinction and CC may be maintained over time^19^. In their within-subject study, two CS+ categories (animals, objects) were used during threat conditioning. Subsequently one of the CS+ categories was used for regular extinction, whereas the other was used for CC. During CC, CS+ exemplars were paired with positively valenced pictures. During a spontaneous recovery task the following day, it was shown that involvement of the vmPFC (amygdala-vmPFC functional connectivity) was stronger for regular extinction compared to CC. In contrast, CS+-specific increases in functional connectivity between the amygdala and the ventral striatum (NAcc) were only observed in the CC condition during a spontaneous recovery task. Both findings are in line with the CC-associated vmPFC deactivation and NAcc activation that we observed and suggest that differences in the neural mechanisms of regular extinction and CC may be maintained during threat retrieval.

CC compared to regular extinction also strengthened item memory for the conditioned category. While both reward and threat conditioning can enhance item recognition for the CS+ category^27,51^, recognition of CS+ exemplars presented during extinction was shown to drop compared to acquisition^31^. In contrast to extinction, within-session CC was previously shown to enhance memory, suggesting that CC has a unique, strengthening effect on memory^29^. In the current study, we replicate this finding, showing strengthened memory after CC compared to extinction. While enhanced recognition of items presented during CC could be mediated by attentional prioritization^52^, CC also retrospectively strengthened memory for items presented during acquisition, suggesting that CC may alter the consolidation of a prior threat conditioning episode. Retroactive enhancement of memory consolidation for related items has previously been shown for conceptually related neutral items presented prior to threat conditioning^32^ and reward conditioning^51^. At a neurobiological level, these findings have been related to the synaptic tagging-and-capture hypothesis postulating that memories for neutral events can be strengthened if they are followed by salient events, due to an initially short-lived synaptic “tag” that allows later events to stabilize the memory^32,53,54^. At a systems level, retroactive memory strengthening has been linked to reverse replay^55^. Specifically, animal research indicates that rewards increase reverse replay^56–58^, and reward-induced reverse replay occurs concurrently with firing of midbrain dopamine neurons^59^. Interestingly, spontaneous replay is also involved in regular extinction, in which unexpected omission of the US drives spontaneous reactivation of activity patterns in the vmPFC. This spontaneous reactivation was shown to be predictive of extinction recall and could be amplified through pharmacological enhancement of dopaminergic activity^60^. Yet while physiological dopaminergic modulation during extinction may be limited to prediction error signals during the early phase^48–50^, dopaminergic modulation may be sustained throughout the MID-based CC task applied in this study. While we did not measure dopaminergic activity directly, activation of the NAcc during reward anticipation is predictive of dopamine release within the NAcc^61–64^. Given the increased stimulus-specific activation of the NAcc in the CC group, it is likely that dopaminergic activity was enhanced during CC compared to regular extinction. The enhanced dopaminergic modulation could strengthen memories through replay^56,65^, or may increase synaptic plasticity directly, potentially explaining enhanced item recognition after CC compared to regular extinction^55,66,67^. In line with these findings, research in humans shows that reward systematically modulates memory for neutral objects in a retroactive manner, with objects closest to the reward being prioritized^55^. It could be that reward-conditioning during CC similarly drives reward-driven reverse replay, enhancing episodic memory for conceptually related items presented during the preceding acquisition task.

Several limitations of the current study are worth considering. First, while the monetary incentive aspect during CC clearly induced positive valence, it also increased physiological arousal, making it difficult to isolate the individual effects of positive valence and reward-induced arousal. While the current results are in line with previous work in CC using low-arousal, positive-valence pictures^29^, we cannot exclude the possibility that the current findings (in part) reflect differences in task engagement between participants due to active instead of passive reward delivery. However, it is questionable whether it is meaningful to tease individual effects of valence and arousal apart since arousal may facilitate reward processing. Indeed, striatal responses to obtained monetary rewards are dependent on salience and are increased when rewards are dependent on active responses compared to passive delivery^68^. Second, although we included a reinstatement procedure in the experiment, neither the Ext nor the CC group showed differential reinstatement. It is worth noting however that reinstatement paradigms in humans may not reliably produce differential reinstatement after extinction^69^. Third, it is important to note that CC/extinction was carried out within minutes after the acquisition phase, and the effects of CC and extinction may differ when carried out after the acquisition memory has been consolidated ^70–73^. Fourth, whole-brain analysis of the CS-specific activation during the spontaneous recovery test in the Ext group did not yield any clusters above threshold, while physiological results indicated spontaneous recovery of differential threat responses. Given that recovered threat responses are often quick to extinguish and fMRI analyses require averaging across multiple trials to achieve sufficient signal-to-noise ratio, threat-evoked neural activity may have been too brief to be detected. Lastly, there was an unequal sex distribution in our sample and the sample size did not allow for the investigation of sex-dependent differences, which should be addressed in future studies.

In conclusion, our findings show that appetitive CC improves the retention of safety memory over standard extinction. Strikingly, in contrast to activation of the vmPFC during extinction, CC was associated with stimulus-specific deactivation of the vmPFC. These findings may inform development of future treatments for fear- and anxiety disorders. While a large body of research focuses on enhancing regular extinction, this study indicates that another promising and potentially longer-lasting approach may be to engage reward-circuits. Although further work is needed, a major advantage of CC-based interventions over extinction-based interventions may be that CC could be more tolerable as it may shift attention away from the experience of fear.

## Materials and Methods

### Participants

Forty-eight healthy right-handed volunteers (15 males, 33 females; age [22.71±0.44]) with no neurological or psychiatric history, and with normal hearing and normal or corrected-to-normal vision completed the study. Exclusion criteria were pregnancy, disorders of the autonomic system, heart conditions, recreational drug use and any contraindications for MRI. Participants provided written informed consent and were paid 55 euros for their participation. Participants in the CC group were able to earn an additional 14 euros. This study was approved by the local ethical review board (METC Oost-Nederland and CMO Radboudumc). Participants were excluded from the threat acquisition, CC/extinction, spontaneous recovery, and reinstatement analyses if there was no evidence for successful threat acquisition (mean CS->CS+ or CS+=CS-). For SCRs this was the case for three participants, for PDR this was the case for two participants. Additional participants were excluded in case of (partially) missing data due to technical failure (no data could be recorded or >50% missing data) or in case participants did not return to the second session. These exclusion criteria were preregistered.

### Design and procedure

This study was a two-day between-subjects experiment carried out in the fMRI scanner (see Figure 1 for an overview of the design). Participants were assigned to either the CC or Ext group according to a predetermined allocation sequence. At the start of each session, two Ag/AgCl electrodes were attached to the medial phalanges of the second and third digit of the left hand, a pulse oximeter was attached to the first digit of the left hand to measure finger pulse and a respiration belt was placed around the abdomen to measure respiration. All measures were taken using a BrainAmp MR system and recorded using the BrainVision Recorder software (Brain Products GmbH, Munich, Germany). The first day consisted of individual adjustment of the electrical shock followed by a single fMRI session that included the following tasks: an object localizer task (17 min), a category threat conditioning task (23 min) and a CC or extinction task (23 min). The second session took place the following day and consisted of three runs: the spontaneous recovery and reinstatement test (12 min), item recognition test (29 min) and the valence-specific response characterization task (17 min).

### Pavlovian conditioning paradigm

Note that CC included an instrumental and not Pavlovian conditioning procedure. This was done because of pragmatic constraints in studies with humans. For example, we cannot food deprive humans to make an appetitive reward truly reinforcing and make participants anticipate the reward. Previous work^51,68^ and our pilot studies indicated that to maximize reward anticipation and evoke conditioned responses, the reward conditioning needed to be instrumental.

The acquisition, counterconditioning, extinction, spontaneous recovery and reinstatement tasks consisted of a categorical differential delay threat conditioning paradigm^27^ with elements of the monetary incentive delay task^30^. Participants viewed trial-unique exemplars of pictures from two categories (animals or objects, see Figure 1). In a counter-balanced manner, exemplars from one category served as CS+ (reinforced) stimuli, while exemplars form the other category served as CS-(unreinforced stimuli). Each trial started with the presentation of a stimulus. After a variable delay of 2.5-4s, a cue appeared to which participants were instructed to respond as quickly as possible with a button press. After the button press, or when a 1s response window had elapsed, the colour of the cue shifted from black to blue. 0.5-1.5s after the response window elapsed, CS+ items presented during the acquisition phase could be reinforced with a shock. During the acquisition phase, 50% of the CS+ pictures were followed by a shock. After 1s, the stimulus was replaced by neutral feedback during the acquisition, extinction, and recovery tasks. During the CC phase, neutral feedback was replaced by monetary feedback. During the CC phase, participants could obtain a €0.50 reward for their quickest responses to the cues presented on top of CS+ stimuli. The response time target was dynamically adjusted to achieve a reward reinforcement rate of approximately 70%. Reward was withheld during the first three CS+ trials during the CC phase to make the transition from the acquisition to the CC phase more gradual. The inter-trial interval (ITI) varied randomly between 8 and 10s. Pictures were presented in a pseudorandom order with no more than 3 consecutive presentations of items from the same category and CC blocks consisted of 40 CS+ and 40 CS-presentations each. The spontaneous recovery block consisted of 15 CS+ and 15 CS+ presentations, and the reinstatement test consisted of 5 CS+ and 5 CS-presentations.

### Item recognition memory test

Participants carried out a surprise recognition memory test compromised of 160 pictures (80 CS+, 80 CS-) shown during the acquisition and CC/extinction phases, as well as 160 category-matched new items (80 CS+, 80 CS-). Participants rated on a 6-point scale whether the picture was ‘definitely old’, ‘probably old’, ‘maybe old’, ‘maybe new’, ‘probably new’, ‘definitely new’.

### Valence-specific response characterization

The valence-specific response characterization task consisted of an adapted version of the conditioning paradigm used during the acquisition phase. Instead of category items, participants were presented with squares in three different colours. One of the stimuli was reinforced with shocks (CS+-shock, 50% reinforcement rate), one stimulus was reinforced with monetary rewards (CS+-reward, approximately 70% reinforcement rate, response time target adjusted dynamically) and the last stimulus was not reinforced (CS-). Each stimulus was presented 40 times in a pseudorandom order with no more than three repetitions of each stimulus. Colours and reinforcement (shocks vs. rewards) were counterbalanced across participants.

### Peripheral stimulation

Electrical shocks were delivered using two Ag/AgCl electrodes attached to the medial phalanges of the second and third digit of the right hand using a MAXTENS 2000 (Bio-Protech) device. Shock intensity varied in 10 intensity steps between 0 to 40 V and 0 to 80 mA. Shock duration was 200 ms. In line with prior threat conditioning protocols, shock intensity was calibrated using an ascending staircase procedure starting with a low voltage setting near a perceptible threshold and increasing to a level deemed “maximally uncomfortable but not painful” by the participant^32,74,75^.

### Arousal and valence ratings

Arousal and valence ratings were acquired using self-assessment manikin scales. The arousal scale ranged from 1 (=extremely calm) to 10 (=extremely excited). The valence scale ranged from 1 (=extremely negative) to 10 (=extremely positive). The valence and arousal ratings were collected for the two categories (animals and tools) after the acquisition phase, after the CC/extinction phase, at the start of day 2 immediately before the spontaneous recovery test and after the reinstatement test. For the stimuli used in the valence-specific response characterization task, valence and arousal ratings were collected immediately after the task.

### SCR pre-processing and analysis

Electrodermal activity data were pre-processed using in-house software; radio frequency (RF) artefacts were removed and a low-pass filter was applied^76,77^. Skin conductance responses (SCR) were automatically scored with additional, blinded, manual supervision using Autonomate^78^. SCR amplitudes (measured in μSiem) were determined for each trial as the maximum response with an onset between 0.5 and 7.5s after stimulus onset and maximum rise time of 14.5s. Shock- and reward-reinforced trials were excluded from analysis. All response amplitudes were square-root transformed and normalized according to each participant’s mean UCS response prior to statistical analysis. The average SCRs were computed per CS-type, task, phase (early, late), and participant.

### PDR pre-processing and analysis

Pupil dilation was measured with a MR-compatible eye-tracker from SensoMotoric Instrument (MEye Track-LR camera unit, SMI, SensoMotoric Instruments) and sampled at a rate of 50 Hz. Data were analysed using in-house software^79^ implemented in Matlab R2018b (MathWorks), based on previously described methods^80^. Eyeblink artifacts were identified and linearly interpolated 100 ms before and 100 ms after each identified blink. Data from scan runs missing 50% time points or more were excluded. After interpolating missing values, time series were band-pass filtered at 0.05 to 5 Hz (by subtracting the mean and dividing by the standard deviation) within each participant and run to account for between-subjects variance in overall pupil size. Event-related pupil diameter responses were calculated by averaging pupil diameter during 3.5 to 7 sec period after stimulus onset, divided by the 1 sec pre-stimulus pupil diameter (−1 to 0 sec). The average PDRs were computed per CS-type, task, phase (early, late), and participant.

### MRI data acquisition

MRI scans were acquired using a Siemens (Erlangen, Germany) 3T MAGNETOM PrismaFit MR scanner equipped with 32-channel transmit-receiver head coil. The manufacturer’s automatic 3D-shimming procedure was performed at the beginning of each experiment. Participants were placed in a light head restraint within the scanner to limit head movements during acquisition. Functional images were acquired with multi-band multi-echo gradient echo-planar (EPI) sequence [51 oblique transverse slices; slice thickness, 2.5mm; TR, 1.5s; flip angle, 75°; echo times, 13.4, 34.8, and 56.2 ms; FOV, 210 x 210 mm2; matrix size 84×84×64, fat suppression]. To account for regional variation in susceptibility-induced signal drop out, voxel-wise weighted sums of all echoes were calculated based on local contrast-to-noise ratio after which echo-series are integrated using PAID weighting^81^. Field maps were acquired (51 oblique transverse slices; slice thickness, 2.5mm; TR, 0.49 s; TE, 4.92 ms and 7.48 ms; flip angle, 60°; FOV, 210 x 210 mm2; matrix size 84×84×64) at the start of each session to allow for correction of distortions due to magnetic field inhomogeneity. A high-resolution structural image (1mm isotropic) was acquired using a T1-weighted 3D magnetization-prepared rapid gradient echo sequence [MP-RAGE; TR, 2300 ms; TE, 3.03 ms; flip angle, 8°; 192 contiguous 1 mm slices; FOV = 256 x 256 mm2].

### fMRI analysis

Anatomical and functional data were pre-processed using fMRIPrep 20.0.6^82^. The complete boilerplate can be found in Supplementary Information 1. In brief, MRI data were pre-processed in standard stereotactic (MNI152) space. Pulse and respiration data were processed offline using in-house software and visually inspected to remove artefacts and correct peak detection, and corrected pulse and respiration data were used for retrospective image-based correction (RETROICORplus) of physiological noise artefacts in BOLD-fMRI data^83^. Identical transformations were applied to all functional images, which were resliced into 2 mm isotropic voxels. After pre-processing in fMRIPrep, functional images were smoothed with a 6 mm FWHM Gaussian kernel (using SPM12; http://www.fil.ion.ucl.ac.uk/spm; Wellcome Department of Imaging Neuroscience, London, UK).

For the acquisition, extinction/cc and spontaneous recovery phases, BOLD responses to CS+, and CS-during the early phase (first half of the trials) and late phase (second half of the trials) were modelled in 4 separate regressors using box-car functions. Additionally, during all these phases, target presentation, button press and shocks were modelled using stick functions, and feedback presentation and breaks were modelled using box-car functions and included as nuisance regressors. For the category localizer, BOLD responses to animals, objects, and phase-scrambled blocks were modelled in 3 separate regressors using box functions. All first-level models also included six movement parameter regressors (3 translations, 3 rotations) derived from rigid-body motion correction, 25 RETROICOR physiological noise regressors, high-pass filtering (1/128 Hz cut-off), and AR(1) serial correlations correction. First-level contrasts were calculated for early and late CS+ and CS-separately for the acquisition, CC/extinction, and spontaneous recovery phases.

For the acquisition and CC/extinction, first-level contrast were entered into a second-level Group (extinction, cc) x CS-type (CS+, CS-) x Phase (early, late) mixed factorial model using the Multilevel and Repeated Measures (MRM) toolbox^84^. For the spontaneous recovery test, BOLD-responses from the early phase were entered into a second-level Group (extinction, cc) x CS-type (CS+, CS-) mixed factorial model. Thresholding was achieved using nonparametric permutation testing (5,000 iterations), with a cluster-setting threshold of p<.001 for whole-brain analysis and familywise error (FWE) correction at p<0.05 at cluster-level for whole-brain analysis and voxel-level for ROI-analysis (Amygdala, Hippocampus, vmPFC, NAcc). Activations are displayed on the single-subject high-resolution T1 volume provided by the Montreal Neurological Institute (MNI).

### Region of interest definition

Based on a priori hypotheses, results for the amygdala, NAcc, hippocampus and the ventromedial prefrontal cortex are corrected for reduced search volumes using small volume. Masks were created using the WFU PickAtlas toolbox^85^ in combination with the Automated Anatomical Labeling atlas^86^ for the bilateral amygdala, bilateral hippocampus and vmPFC (Frontal_Med_orb_L&R and Rectus L&R). The *IBASPM 71* anatomical atlas toolbox^87^ as used to create a mask for the bilateral NAcc.

### Statistical testing

Statistical analyses of behavioural and physiological variables were performed in SPSS (IBM SPSS Statistics Inc.). Dependent measures were submitted to repeated measure ANOVAs and statistics were Greenhouse-Geisser or Huyn-Feldt corrected for non-sphericity when appropriate. Repeated measures ANOVAs were applied to capture early versus late phases of acquisition and CC/extinction, as well as to compare late CC/extinction (last 2 trials) compared to early spontaneous recovery (first 2 trials). We refrained from trial-by-trial analyses in a relatively small sample as this would have cost too many degrees of freedom and is not expected to provide more information. Significant findings from ANOVAs were followed-up by paired- and independent samples t-tests and Bonferroni-adjusted where applicable. We report partial eta-square as measure of effect size. Means ± s.e.m are provided where relevant unless otherwise indicated.

### Deviations from the pre-registration

The preregistration for this project can be found on OSF (https://osf.io/fbz6n). We pre-registered to sample SCRS in a 0.75 and 3.15 s window after stimulus onset. However, visual inspection of SCR responses during the acquisition phase indicated that response latencies shifted towards the late phase of the trial. We therefore opted to use a longer window (0.5s to 7.5s for stimulus onset) and exclude reinforced trials. The pre-registration erroneously stated that pupil-dilation data would be z-scored and later divided by the pre-stimulus average. PDR data were not z-scored but were only normalized to a 1-sec pre-stimulus baseline. In line with the SCR data, response onset latencies were later than expected. Based on visual inspection of the data from the acquisition phase, we decided to use a window around the expected shock onset: 3.5-7s after stimulus onset. Reinforced trials were excluded. Results for SCR, retrospective reinforcement estimations and the reinstatement test can be found in the **Supplementary Information**. Due to an error in the scripts for the item recognition test, trial-by-trial data were not recorded for the first 12 participants. Therefore, analysis of the memory data focused on averaged data for the early and late phase of acquisition and CC/extinction, leaving out planned change point analyses on bins of 4 trials.

While we planned to extract a vmPFC mask for ROI analysis based on a [CS-> CS+ shock] contrast of BOLD responses during the valence-specific response characterization task to identify “extinction regions”, this did not yield ventromedial prefrontal clusters that survived correction. Instead, in line with our other ROIs, we opted to create a mask based on the AAL atlas. Due to time constraints, native-space and functional connectivity analyses were not carried out for this manuscript.

## Data Availability

Pseudonymized data generated in this study are available upon request from the Radboud Data Repository at https://data.ru.nl/. Raw MR images are not publicly available due to privacy or ethical restrictions.

## Acknowledgement

This work was supported by the European Research Council (ERC-2015-CoG 682591).

## Author contributions

M.C.H., J.E.D., J.H., M.J.A.G.H. and E.H. designed the study. M.C.H. and J.V. implemented and conducted the experiment. M.C.H., L.W. and J.V. analysed the results with support of E.H.. L.W. and M.C.H. wrote the paper with contributions from all authors.

## Supplementary Information

### Valence-specific response characterization

At the end of the experiment, participants underwent a simplified version of the main experimental task, in which category exemplars were replaced by colored squares. This task was used to investigate to what extent skin conductance responses (SCRs) and pupil dilation responses (PDRs) can be used to disentangle anticipation of shock and reward. Participants viewed three different coloured squares and learned that one colour was associated with shocks (CS+S), one colour with rewards (CS+R) and one colour served as CS-. The trial structure was otherwise identical to comparable trials from the acquisition and CC phases. At the end of the task, participants were asked to rate the three stimuli on valence and arousal self-assessment manikin scales (Bradley & Lang, 1994).

During this valence-specific response characterization task, we observed habituation in SCRs over the course of the task (CS-type (CS+ S, CS+ R, CS-) x Phase (early, late) x Group (CC, Ext) rmANOVA, main effect phase: F(_1,38_)=13.921, p=0.001, η²=0.268) and different SCR magnitudes for the three different CS-types (main effect CS-type (CS+S, CS+R, CS-): F(_2,76_)=78.460, p<0.001, η²=0.674). In addition, habituation depended on CS-type (CS-type x Phase interaction: F(_2,76_)=6.825, p=0.002, η²=0.152). During the early phase, SCRs in response to the CS+R and the CS-were not distinguishable (t(40)=0.115, p=0.909, CS+R: 0.32±0.03, CS-:0.32 ±0.03), while during the late phase, SCRs to the CS+R were larger than the CS-(t(40)=4.993, p<0.001, CS+R: 0.29±0.03, CS-:0.19±0.02). SCRs to the CS+S were consistently larger than SCRs to the CS+R (early: t(41)=9.345, CS+S: 0.62±0.04, p<0.001, late: t(40)=5.952, p<0.001, CS+S: 0.56±0.04) and the CS-(early: t(40)=10.020, p<0.001, late: t(4)=10.122, p<0.001). Thus, anticipation of aversive reinforcement (CS+S) led to increased SCRs compared to anticipation of reward (CS+R) and CS-presentation throughout the task. Due to the fact that SCRs performed less well in differentiation between CS+R and CS-, we focused our analyses on PDRs, but report SCR results here as well.

We also observed CS-type dependent differences in PDRs (CS-type (CS+ S, CS+ R, CS-) x Phase (early, late) x Group (CC, Ext) rmANOVA, main effect CS-type (CS+S, CS+R, CS-): F(_2,68_=19.783, p<0.001, η²=0.368). In comparison to the neutral CS-, both the shock-reinforced CS+ (CS+S) and reward-reinforced CS+ (CS+R) evoked larger PDRs (Supplementary Figure 1A, t(36)=7.071, p<0.001 and t(26)=4.900, p<0.001 respectively, CS+S: 1.05±0.03, CS+R: 1.04±0.04, CS-: 1.01±0.02). However, reward- and shock anticipation-induced PDRs did not differ statistically (t(36)=1.146, p=0.259). While both shock anticipation and reward anticipation led to similar increases in PDRs as compared to the neutral condition, valence and arousal ratings indicated that participants experienced shock and reward trials differently. Specifically, the CS+R was rated more positive than the CS-(t(47)=9.046, p<0.001, CS+R: 7.79±0.14, CS-: 5.96±0.16, Supplementary Figure 1C), while the CS+S was rated less positive than the CS-(t(47)=-10.337, p<0.001, CS+S: 2.96±0.25). Participants reported increased arousal to both the CS+S and CS+R as compared to the CS-(t(47)=4.666, p<0.001 and t(47)=8.897, p<0.001 respectively, CS+S: 5.42±0.35, CS+R: 6.31±0.21, CS-: 3.33±0.30, Supplementary Figure 1B). While it was not possible to distinguish PDRs to the CS+S and CS+R, explicit ratings of arousal were marginally increased for the CS+R as compared to the CS+S (t(47)=-2.100, p=0.041). In conclusion, the response characterization task showed that while anticipation of reward and shock both generate increased PDRs as compared to the CS-, distinct retrospective valence ratings show the expected directions.

**Supplementary Figure 1.**
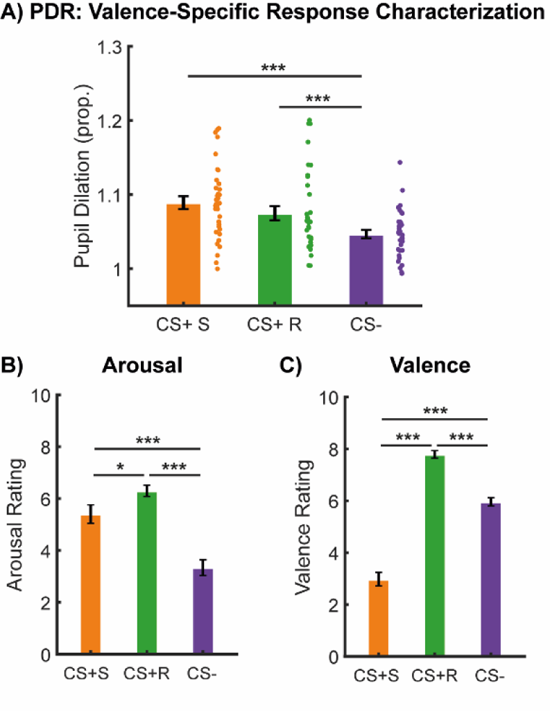
Pupil dilation responses (PDRs), explicit arousal and valence rating for the different CSs presented during the valence-specific response characterisation task. (A) PDRs to the shock reinforced (CS+S), reward reinforced (CS+R) and CS-stimuli, averaged across the task and all participants. PDRs were increased for the CS+S and CS+R as compared to the CS-(B) Explicit ratings of arousal and (C) valence provided immediately after the task. Explicit ratings of arousal for the CS+S exceeded ratings for the CS-, and the CS+R was rated higher in arousal than the CS+S. Valence ratings (1=extremely negative, 10=extremely positive) for the CS+R were more positive than ratings for the CS-, while ratings for the CS+S were more negative than for the CS-and CS+R. Error bars represent ± standard error of the mean *=p<0.05, ***=p<0.001

### Threat acquisition

#### Physiological and behavioural evidence for acquisition of conditioned threat responses

Participants pre-assigned to the CC and Ext groups underwent an identical threat acquisition procedure. To verify that participants pre-assigned to both groups acquired conditioned threat responses of comparable strength, we compared PDRs, explicit valence and arousal ratings, and response times between groups. During the acquisition task, participants pre-assigned to both groups showed stable and comparable differential conditioned threat responses as measured by PDRs (Supplementary Figure 2A, CS-type (CS+, CS-) x Phase (Early, Late) x Group (CC, Ext) rmANOVA, main effect CS-type: F(_1,37_)=41.172, p<0.001, η²=0.533, other main effects and interactions: all p’s>0.2). Both groups also acquired comparable differential SCRs (main effect CS-type: F(_1,42_)=58.633, p<0.001, η²=0.583), although SCRs showed habituation over the course of the task (main effect phase: F(_1,42_)=66.907, p<0.001, η²=0.614, all other p’s>0.3). Thus, both SCRs and PDRs demonstrated comparable acquisition of conditioned threat responses between groups.

Successful threat acquisition was further confirmed by valence and arousal ratings for the CS+ and CS-categories at the end of the acquisition task. Arousal ratings for the CS+ category exceeded arousal ratings for the CS-category (Supplementary Figure 2B, CS-type (CS+, CS-) x Group (CC, Ext) rmANOVA, main effect CS-type: F(_1,44_)=27.573, p<0.001, η²=0.385), and did not differ between groups (all p’s>0.2). Similarly, the CS+ category was given lower valence (less positive) ratings than the CS-category (Supplementary Figure 2C, CS-type (CS+, CS-) x Group (CC, Ext) rmANOVA, main effect CS-type: F(_1,44_)=12.626, p<0.001, η²=0.223). Although there was no main effect of group on valence ratings (p>0.7), the effect of CS-category unexpectedly differed between the CC and Ext group (CS-type x Group interaction: F(_1,44_)=4.512, p=0.039, η²=0.093), due to more positive ratings to the CS-category in the Ext group (CC: 5.8±0.4, Ext: 6.9±0.3, t(44)=2.156, p=0.037). Nevertheless, valence ratings for the CS+ category were comparable between groups (CC: 5.1±0.5, Ext: 4.2±0.4, p>0.1), suggesting that the strength of the acquired threat responses is likely similar between groups.

**Supplementary Figure 2.**
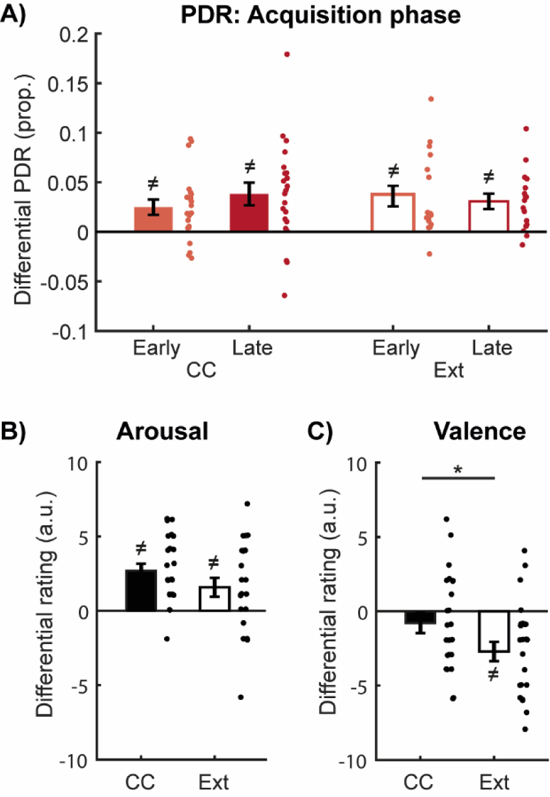
Differential PDRs during acquisition and explicit ratings of arousal and valence provided after acquisition. (A) Differential PDRs for the early (light red) and late (dark red) phase of the acquisition task, (B) arousal and (C) valence ratings, displayed separately for participants assigned to the counterconditioning (CC, solid bars) and extinction (EXT, open bars) groups. Both groups showed comparable differential PDRs and arousal ratings during the acquisition task. Participants in both groups showed negative differential valence ratings (stronger negative valence for CS+ vs. CS-), although this effect was stronger in the Ext group. Error bars represent ± standard error of the mean. *, p<0.05. ≠. Significantly different from 0

To keep all experimental tasks similar between groups, participants in both groups were asked to respond to targets that were superimposed on the stimuli as quickly as possible. To verify that both groups performed similarly on this task, we compared response times for the different stimuli between the groups. During the acquisition task, participants responded faster to targets in CS+ trials compared to CS-trials (CS-type (CS+, CS-) x Group (CC, Ext) rmANOVA, main effect CS-type: F(_1,45_)=10.839, p=0.002, η²=0.194), with no differences between groups (all p’s>0.058).

After the acquisition task, participants in both groups reported higher estimated reinforcement rates for the CS+ category as compared to the CS-category (CS-type (CS+, CS-) x Group rmANOVA, main effect CS-type: F(_1,45_)=82.176, p<0.001, η²=0.646). The reported reinforcement rates did not differ between groups (all p’s>0.3).

#### Brain activation supports successful acquisition of conditioned threat responses

The acquisition of conditioned fear on the first day reliably activated networks associated with fear conditioning. Whole-brain analysis identified regions that were more responsive to the CS+ versus the CS-category (Supplementary Figure 3, see Supplementary Table 1 for a complete overview of findings). We observed differential BOLD responses in a large number of brain areas, including the bilateral insula, posterior and anterior cingulate, thalamus, precuneus (undirected test, cluster size = 425400 mm^3^, p<0.001, whole-brain FWE-corrected) and the bilateral amygdala (right cluster size = 1088 mm^3^, p<0.001, FWE-SVC, left cluster size = 736 mm^3^, p<0.001, FWE-SVC).

**Supplementary Figure 3.**
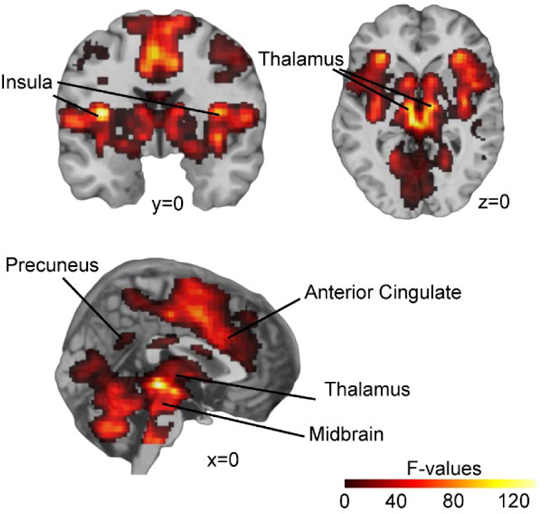
Differential threat responses during acquisition revealed CS-specific activation of clusters encompassing a range of regions including the bilateral insula, thalamus, precuneus, anterior cingulate and midbrain. Group F-image of the effect of CS type, thresholded at cluster-level FWE-corrected p<0.05, cluster-forming threshold p=0.001, displayed on the single-subject high-resolution T1 volume provided by the Montreal Neurological Institute (MNI).

**Supplementary Table 1.**
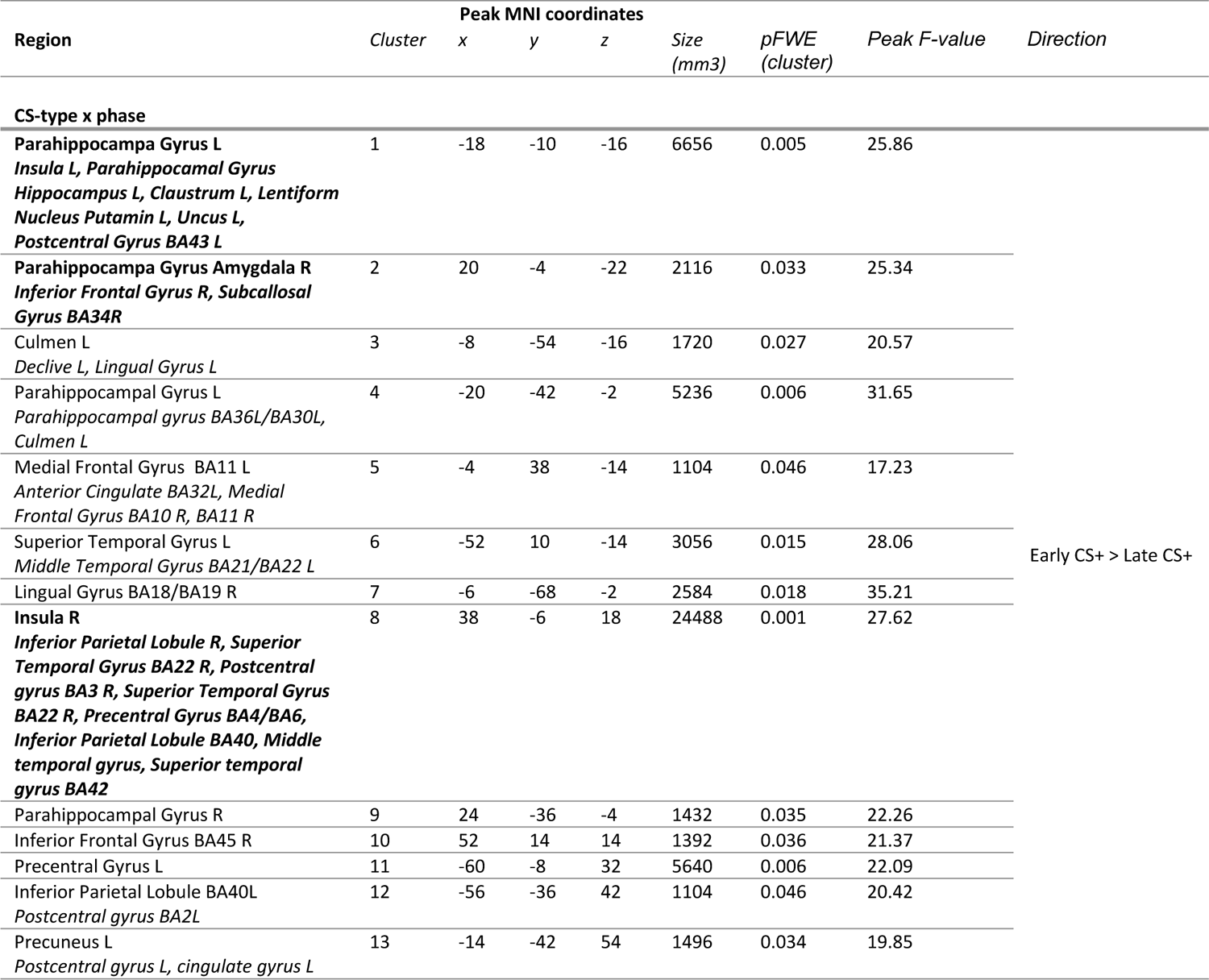
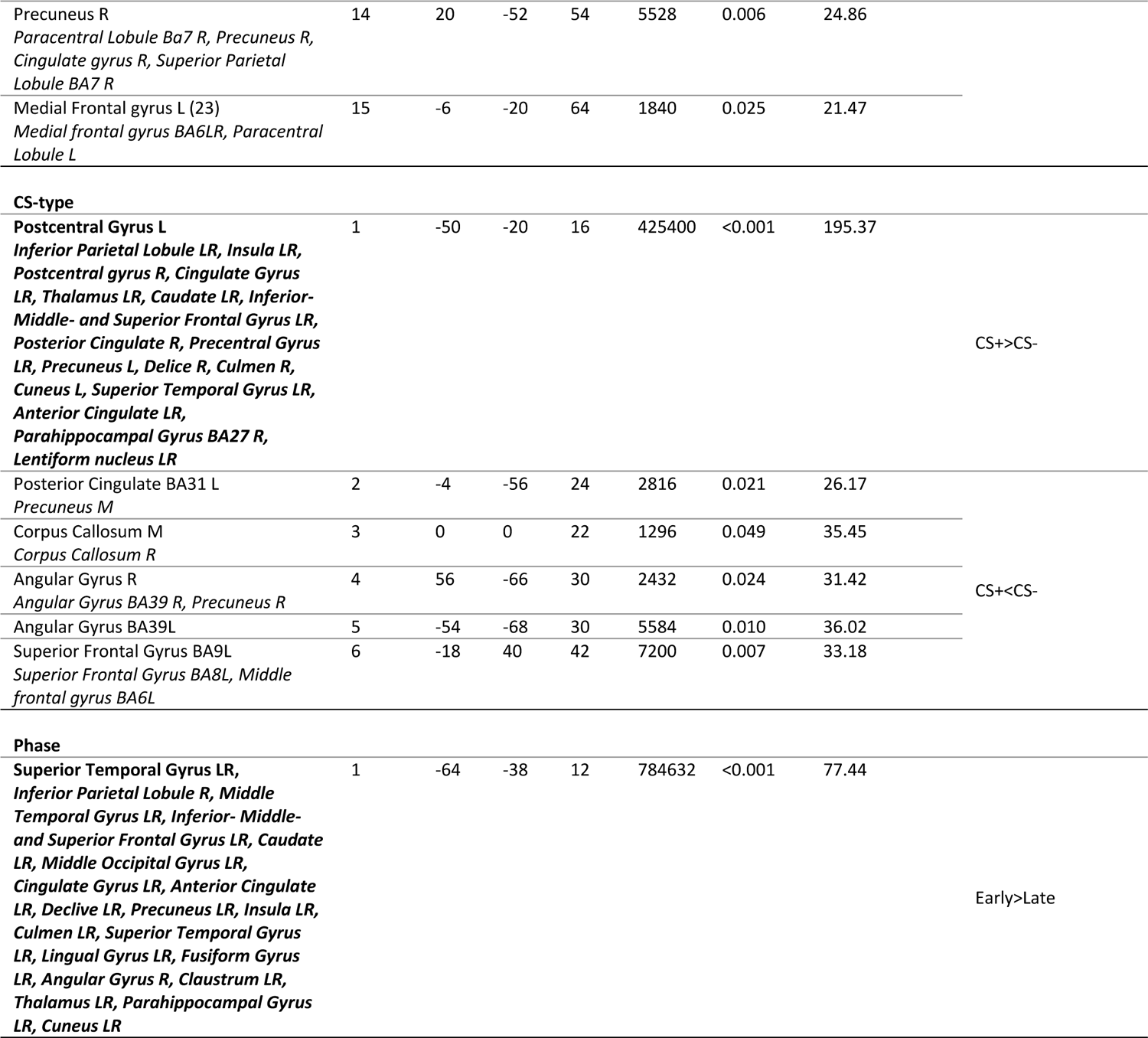
Whole-brain main effects of group (CC, Ext), CS type (CS+, CS-) and phase (early, late) and interactions, during the acquisition task. Cluster-forming threshold p=0.001, FWE-corrected at p<0.05, clusters were labelled using the Talairach Deamon atlas and the AAL atlas for ROIs. For each cluster, the peak voxel coordinates (MNI space) and regions are reported, and additional regions contained within the cluster are added in italics.

### Counterconditioning and extinction

#### Skin conductance responses

Differential SCRs were still apparent during the CC/extinction task (CS-type (CS+, CS-) x Phase (Early, Late) x Group (CC, Ext) rmANOVA, main effect CS-type: F(_1,40_)=17.609, p<0.001, η²=0.306). To verify that successful extinction was reached by the end of the task, we explored SCRs in the late phase separately, but found that differential SCRs were still apparent in that phase (F(_1,41_)=12.166, p=0.001, η²=0.229). Finally, we explored whether the last two trials of the extinction task showed evidence of residual differential SCRs. In the last two trials of extinction, across both groups, there was no evidence for differential SCRs (all p’s>0.2). Thus, while differential SCRs persisted during the late phase of the extinction task, differential responses were no longer apparent in the last two trials. Throughout the CC/extinction, there was no evidence for different SCRs between groups, suggesting that participants in both groups underwent comparable but slow extinction of differential SCRs.

#### Overlapping stimulus-specific activation during counterconditioning and extinction

A number of clusters showed comparable stimulus-specific activations during CC and extinction (Supplementary Table 2).

**Supplementary Table 2.**
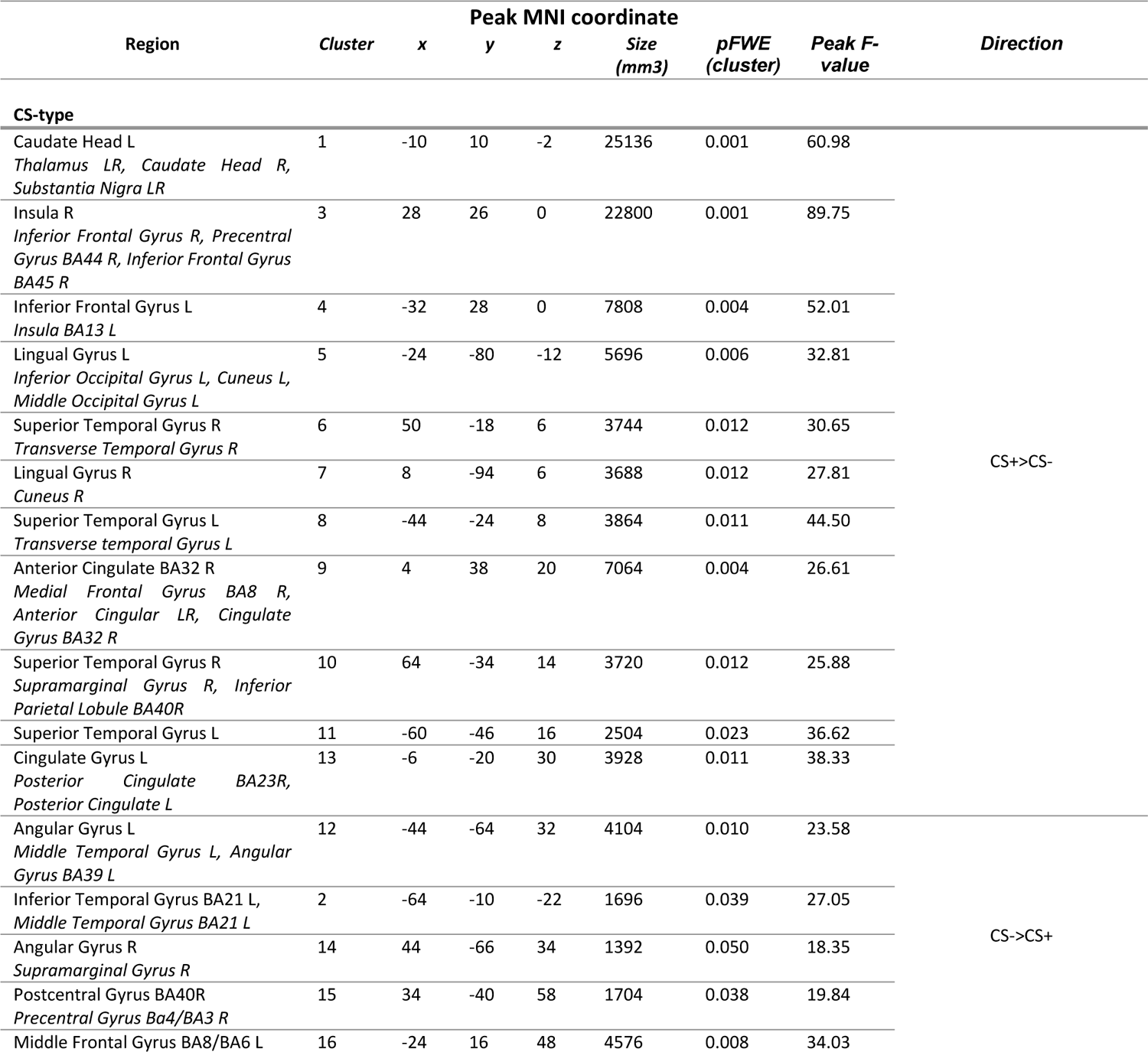
Whole-brain main effect of CS-type during the CC/extinction task. Cluster-forming threshold p=0.001, FWE-corrected at p<0.05, clusters were labelled using the Talairach Daemon atlas and the AAL atlas for ROIs. For each cluster, the peak voxel coordinates (MNI space) and regions are reported, and additional regions contained within the cluster are added in italics.

### Spontaneous recovery test

#### Skin conductance responses

To investigate whether CC can prevent spontaneous recovery of differential SCRs, SCRs during the last two trials of extinction and the first two trials of the spontaneous recovery test were submitted to a CS-type (CS+, CS-) x Phase (last 2 trials of the CC/extinction phase, first two trials of the spontaneous recovery test) x Group (CC, Ext) rmANOVA. SCRs showed a generalized increase from the last two trials of extinction to the first two trials of the spontaneous recovery test (main effect phase: F(1,38)=32.392, p<0.001, η²=0.460)). There was evidence for differential SCRs across both phases (main effect CS-type: (F(1,38)=9.560, p=0.004, η²=0.201), as CS+ stimuli evoked higher SCRs than CS-stimuli (t(43)=2.518,p=0.016, CS+:0.41±0.03, CS-:0.35±0.03), yet we did not find evidence for CS+-specific spontaneous recovery or effect of group (all p’s>0.4). Thus, although there was a generalized increase in responding from the end of extinction to the start of the spontaneous recovery test, SCRs did not show differential recovery and were comparable between groups.

#### Brain activation

During the spontaneous recovery test, CS-specific activation differed between groups in the inferior temporal gyrus (cluster size = 2008 mm^3^, p=0.020, FWE-corrected, Supplementary Figure 4A) and the inferior frontal gyrus (cluster size = 1920 mm^3^, p=0.022, FWE-corrected, Supplementary Figure 4B). Separate analysis of the spontaneous recovery phase within each group did not reveal any suprathreshold clusters in the Ext group, while a number of clusters showed stimulus-specific activation in the CC group. Specifically, the CC group showed stimulus-specific activation in the bilateral fusiform gyri, superior parietal lobes and inferior frontal gyri, and in the right thalamus, caudate, middle frontal gyrus, and angular gyrus (see Supplementary Table 3). A priori defined regions of interest (ROIs) during the spontaneous recovery task were submitted to a Group (CC, Ext) x CS-type (CS+, CS-) x Phase (early, late) rmANOVA but did not reveal any effects.

**Supplementary Figure 4.**
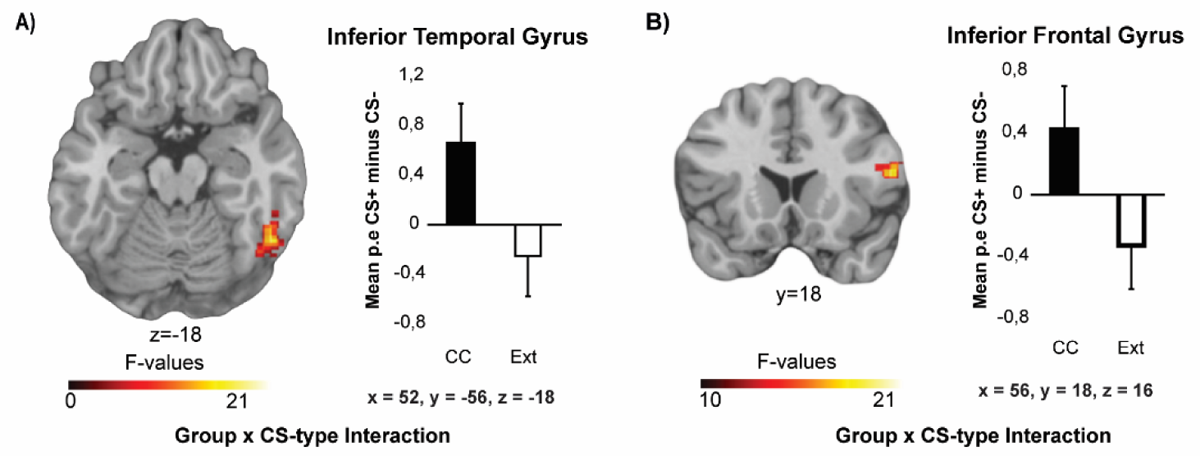
During the spontaneous recovery test, stimulus type-specific activation of the inferior temporal and frontal gyri differed between groups. The inferior temporal Gyrus (A) and Inferior frontal gyrus (B) show increased CS+-specific activation in the CC group as compared to the Ext group. Group F-images thresholded at FWE-corrected p<0.05, cluster-forming threshold p=0.001, displayed on the single*-*subject high-resolution T1 volume provided by the Montreal Neurological Institute (MNI) and parameters estimates from peak voxels.

**Supplementary Table 3.**
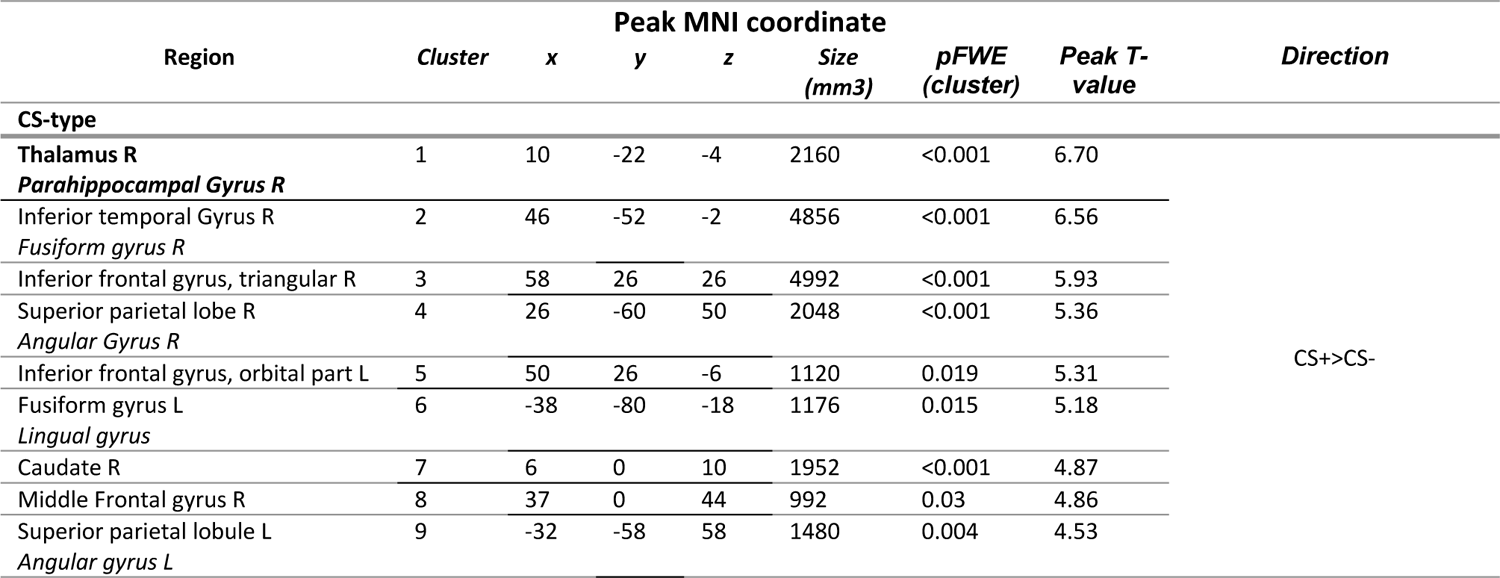
Peak voxel coordinates and statistics of activations during the spontaneous recovery phase in the CC group. Clusters were labelled using the AAL atlas. For each cluster, the peak voxel coordinates and regions are reported, and additional regions contained within the cluster are added in italics. Clusters are whole-brain FWE-corrected at p<0.05.

#### Reinstatement test

SCRs showed a generalized increase from the spontaneous recovery phase to the reinstatement test (CS-type (CS+, CS-) x Group (CC, Ext) x Phase (spontaneous recovery test, reinstatement test) rmANOVA, main effect Phase: F(_1,39_)=25.758, p<0.001, η²=0.398, last two trials of spontaneous recovery: 0.22±0.04, first two trials of reinstatement: 0.38±0.03). Across the last two trials of the spontaneous recovery test and the first two trials of the reinstatement test, differential SCRs differ between the counterconditioning and extinction group (interaction effect of stimulus type and group: F(_1,39_)=4.967, p=0.032, η²=0.113). Yet, there is no evidence for differential reinstatement between groups (no CS-type x Phase x Group interaction, p=0.218). Moreover, mean SCRs to CS+ and CS-stimuli do not differ within either group (all p’s>0.12).

PDRs showed a generalized decrease from the spontaneous recovery phase to the reinstatement test (CS-type (CS+, CS-) x Group (CC, Ext) x Phase (spontaneous recovery test, reinstatement test) rmANOVA, main effect of Phase (F(_1,29_)=9.104, p=0.005, η²=239)). Mean PDRs decreased from spontaneous recovery to reinstatement (t(30)=3.063, p=0.005, last two trials of spontaneous recovery: 1.04±0.01, first two trials of reinstatement: 1.01±0.01). Given that we did not observe successful reinstatement in either group, our reinstatement test does not inform us about whether CC can lead to a more persistent attenuation of fear as compared to classic extinction.

#### CS+-specific enhancement of recognition memory depends on CS+ category

Corrected recognition scores (pHits – pFA) were subjected to a task (acquisition, CC/extinction task) x CS-type (CS+, CS-) x Group (CC, Ext) rmANOVA, including CS+-category (animals, tools) as covariate. Although the effect of CS-type differed depending on the category used as CS+ (CS-type x CS+-category interaction: F(_1,42_)=19.400, p<0.001, η²=0.316) and task (CS-type x CS+-category x task interaction: F(_1,43_)=5.375, p=0.025, η²=0.113), showing a stronger effect for tools as CS+, this did not differ between our experimental groups (CS-type x CS+-category x Group interaction: F(1,41)=0.050, p=0.824; CS-type x task x CS+-category x Group interaction: F(_1,41_)=0.005, p=0.946).

To further investigate to what extent CC retroactively affected memory for items presented during the acquisition task, we examined recognition of items presented during acquisition and the CC/extinction tasks separately. Retrospective memory enhancement for the CS+ items compared to CS-items differed depending on the CS+ category during both the acquisition task (CS-type x CS+ category interaction: F(_1,42_)=29.730, p<0.001, η²=0.414, CS+ category main effect: F(_1,42_)=5.346, p=0.026, η²=0.113) and the CC/extinction task (CS-type x CS+ category interaction: F(_1,42_)=5.121, p=0.029, η²=0.109), showing a stronger effect for tools as CS+. Importantly, this effect was comparable between groups (CS-type x CS+ category x Group interaction: acquisition task F(_1,41_)=0.016, p=0.806; CC/extinction task F(_1,41_)=0.019, p=0.890).

